# A rapid and efficient red-light-activated Cre recombinase system for genome engineering in mammalian cells and transgenic mice

**DOI:** 10.1101/2025.03.16.643490

**Authors:** Yang Zhou, Yu Wei, Jianli Yin, Deqiang Kong, Wenjun Li, Xinyi Wang, Yining Yao, Qin Huang, Lei Li, Mengyao Liu, Longliang Qiao, Huiying Li, Junwei Zhao, Tao P Zhong, Dali Li, Liting Duan, Ningzi Guan, Haifeng Ye

**Affiliations:** Wuhu Hospital, Health Science Center, East China Normal University, Wuhu 241001, China; Shanghai Frontiers Science Center of Genome Editing and Cell Therapy, Biomedical Synthetic Biology Research Center, Shanghai Key Laboratory of Regulatory Biology, Institute of Biomedical Sciences and School of Life Sciences, East China Normal University, Shanghai 200241, China; Yantai Institute of Coastal Zone Research, Chinese Academy of Sciences, Yantai 264003, China; School of Chemistry and Molecular Engineering, East China Normal University, Shanghai 200241, China; Chongqing Key Laboratory of Precision Optics, Chongqing Institute of East China Normal University, Chongqing 401120, China; Department of Biomedical Engineering, The Chinese University of Hong Kong, Sha Tin, Hong Kong SAR 999077, China; Joint Center for Translational Medicine, School of Life Science, East China Normal University and Fengxian District Central Hospital, Shanghai 201499, China; Beijing Life Science Academy, Yingcai South 1st Street, Future Science City South District, Beiqijia Town, Changping District, Beijing 102209, China

## Abstract

The Cre-loxP recombination system enables precise genome engineering; however, existing photoactivatable Cre tools suffer from several limitations, including low DNA recombination efficiency, background activation, slow activation kinetics, and poor tissue penetration. Here, we present REDMAPCre, a red-light-controlled split-Cre system based on the ΔPhyA/FHY1 interaction. REDMAPCre enables rapid activation (1-second illumination) and achieves an 85-fold increase in recombination efficiency. We demonstrate its efficient regulation of DNA recombination in mammalian cells and mice, as well as its compatibility with other inducible recombinase systems for Boolean logic-gated DNA recombination. Using a single-vector adeno-associated virus (AAV) delivery system, we successfully induced REDMAPCre-mediated DNA recombination in mice. Furthermore, we generated a REDMAPCre transgenic mouse line and validated its efficient, light-dependent recombination across multiple organs. To explore its functional applications, REDMAPCre transgenic mice were crossed with the relative Cre-dependent reporter mice, enabling optogenetic induction of insulin resistance and hepatic lipid accumulation via Cre-dependent overexpression of ubiquitin-like with PHD and ring finger domains 1 (UHRF1), as well as targeted cell ablation through diphtheria toxin fragment A (DTA) expression. Collectively, REDMAPCre provides a powerful tool for achieving remote control of recombination and facilitating functional genetic studies in living systems.

## Introduction

Cre recombinase is a site-specific recombinase of the integrase family capable of catalyzing directional DNA recombination between *loxP* pairs^1,2^. Cre-*loxP*-mediated recombination system offers advantages because Cre recombinase does not generate DNA double-strand breaks, which are toxic to the target cell and can trigger undesirable point mutations and structural variations^3,4^. Therefore, recombinase has become the prime tool in various genetic engineering strategies and is wildly employed to generate conditional gene knockout (KO) mice^2,5,6^. A conditional DNA recombination system capable of spatiotemporal genomic manipulation has enabled many advancements in various biological and biomedical research areas^7–10^. Several chemically induced Cre-*loxP* recombination systems have developed to enable temporal control of DNA recombination^11–14^. However, these systems have several limitations. Since chemical inducers act systemically, they lack the ability to achieve precise spatial control, making tissue-specific genome manipulation challenging^15–17^.

In contrast, optogenetic approaches offer spatiotemporal precision, allowing for localized and reversible activation with high temporal resolution^18,19^. Light-based systems eliminate concerns about drug metabolism, toxicity, and systemic diffusion, making them a more precise and controllable alternative for genome engineering^20^. Recent developments of photoactivatable Cre-*loxP* recombination systems have provided spatial and temporal regulation of DNA recombination^21–23^. However, the limited penetration depth of blue/violet light wavelengths significantly restricts their application scope, particularly in transgenic mouse models^24–27^. For instance, the ePA-Cre system shows effective gene recombination only at superficial layers of up to 1 millimeter in targeted organs^22^. Deeper tissue recombination requires the use of implantable optical fiber techniques. Their widespread adoption in research is further prevented by the risk of unintended background recombination activated by ambient light^22,28,29^. Recent study introduced a doxycycline– and violet light-inducible Cre recombinase and resolved unintended light activation, but non-invasive activation in deep tissues remains unfeasible^29^. Notably, the additional introduction of the doxycycline-inducible genetic element increased the payload burden on the optogenetic recombinase system.

To address these limitations, recent efforts have focused on utilizing longer-wavelength light, which offers improved tissue penetration in vivo^30–32^. Red or far-red light-activated Cre systems enable efficient, fiber-free DNA recombination through non-invasive illumination. However, these systems typically require hours of continuous illumination to achieve recombination^31,32^. Furthermore, persistent concerns about unintended recombination due to ambient light exposure, along with the necessity for prolonged illumination, likely account for the absence of transgenic mouse models incorporating red light-inducible Cre recombinase to date.

An ideal photoactivatable recombinase should exhibit high spatiotemporal precision, minimal background activity, and deep-tissue penetrability. Here, we introduce REDMAP_Cre_, a red-light-controlled split-Cre recombinase system engineered to overcome the limitations of existing optogenetic tools. Build upon our previously developed ΔPhyA/FHY1 heterodimerization platform, which responds to 660 nm light^33^, REDMAP_Cre_ utilizes phycocyanobilin (PCB) as a covalently bound chromophore for photoswitching, ensuring negligible background activity in the absence of both PCB and illumination. Upon red-light exposure, REDMAP_Cre_ enables rapid recombination activation within seconds.

For in vivo applications, we delivered REDMAP_Cre_ into liver and muscle tissues using adeno-associated virus (AAV) vectors, achieving successful DNA recombination upon red-light illumination. Furthermore, we established a REDMAP_Cre_ transgenic mouse line, which facilitates precise, light-dependent DNA recombination. By crossing this line with Cre-dependent reporter or functional gene-modified mice, we generated offspring that enable red-light-controlled gene activation or deletion in a tissue-specific manner. This was exemplified by the optogenetic induction of insulin resistance and hepatic lipid deposition via Cre-dependent ubiquitin-like with PHD and ring finger domains 1 (UHRF1) overexpression, as well as targeted cell ablation through diphtheria toxin fragment A (DTA) activation. Collectively, these advancements provide a versatile and robust toolkit for mammalian cell and murine studies, enabling rapid, low-background, and remotely controlled DNA recombination with high spatiotemporal resolution.

## Results

### Design of REDMAP_Cre_ and optimization for enhanced DNA recombination activity

Seeking a highly efficient photoactivatable Cre recombinase, we designed REDMAP_Cre_, which is based on the light-gated reassembly of split-Cre fragments driven by the REDMAP system (a chromophore-dependent red-light-inducible ΔPhyA/FHY1 dimerization system^33^). ΔPhyA is fused with C-terminus of Cre (CreC), and FHY1 with N-terminus of Cre (CreN); the chromophore phycocyanobilin (PCB) binds covalently to ΔPhyA, making it responsive to 660 nm red light. Under red light and PCB, ΔPhyA quickly binds to FHY1, reconstituting the split-Cre fragments that excise DNA flanked by two *loxP* sites **(Fig. 1a)**.

**Figure 1.**
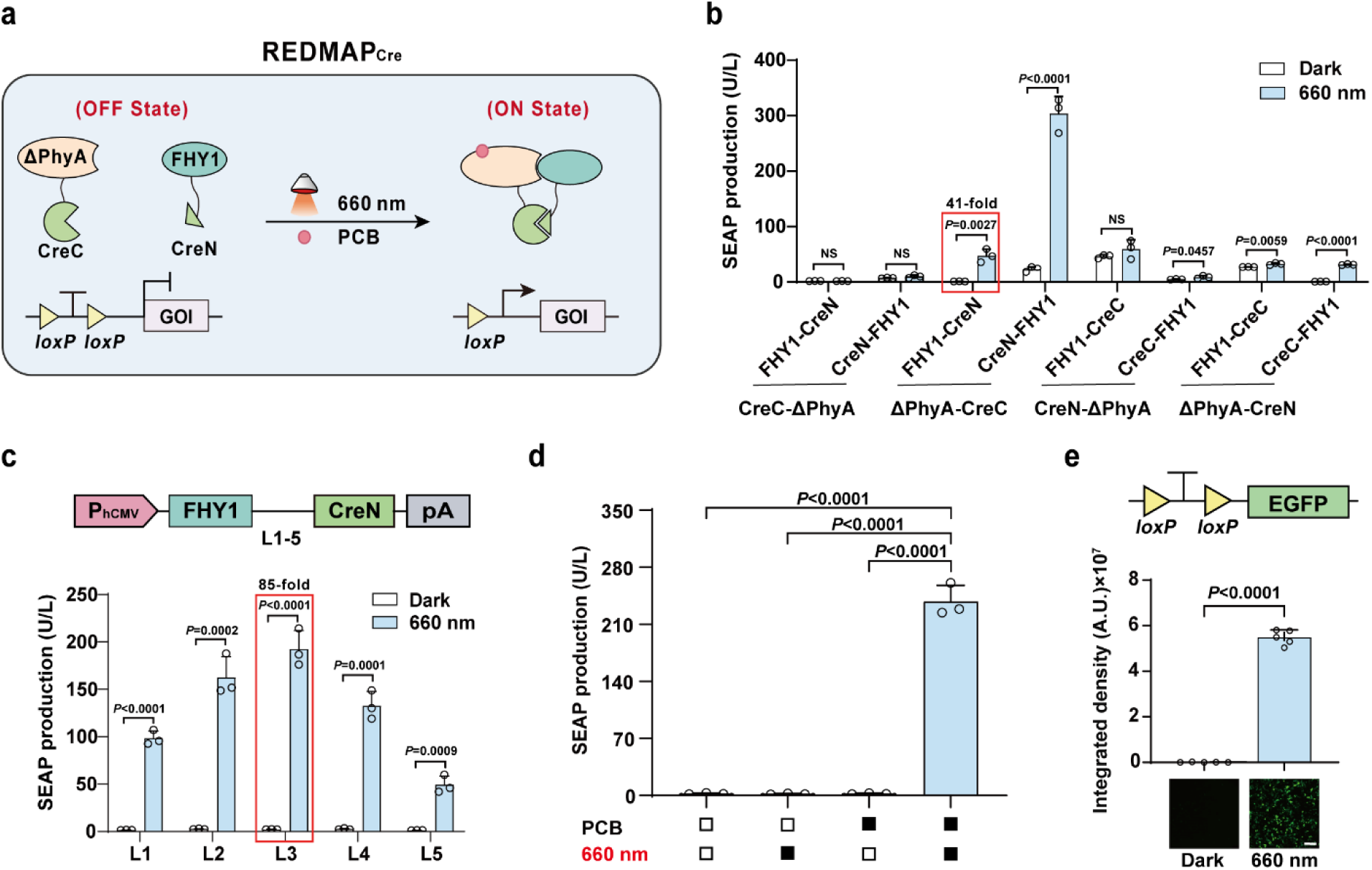
| Design and optimization of REDMAP_Cre_. **a**, Schematic for the photoactivatable Cre system (REDMAP_Cre_). Cre recombinase was split into two fragments: CreN59 (residues 1-59) fused to FHY1 and CreC60 (residues 60-343) fused to ΔPhyA, with the expression of each driven by a constitutive promoter. In the presence of the photosensitive pigment PCB, ΔPhyA and FHY1 dimerize upon 660 nm light illumination, resulting in the restoration of Cre recombinase catalytic activity, enabling excision of DNA sequences (STOP) flanked by *loxP* sites. **b**, Screening the indicated combinations of ΔPhyA/FHY1 domains and split Cre (59/60) fragments. HEK-293T cells (6×10^4^) co-transfected with the various fusion protein expression vectors and the Cre-dependent SEAP reporter (pGY125, P_hCMV_-*loxP*-STOP-*loxP*-SEAP-pA), were supplied with PCB (5 μM) and then illuminated with red light (660 nm, 1 mW cm^-2^) for 48 hours. SEAP production in the culture supernatant was quantified after illumination. **c**, Optimization for recombination efficiency using the different linkers (L1 to L5) between the FHY1 and CreN domains. HEK-293T cells (6×10^4^) were co-transfected with ΔPhyA-L5-CreC expression vector (pYZ208), pGY125, and FHY1-CreN fusion protein expression vector with different linkers (L1 to L5), with treatment as described in **b**. SEAP production was quantified 48 hours after illumination. **d**, REDMAP_Cre_-catalyzed DNA recombination is dependent on red light and on PCB. HEK-293T cells (6×10^4^) co-transfected with ΔPhyA-L5-CreC (pYZ208), FHY1-L3-CreN (pYZ247), and pGY125, were supplied with or without PCB, followed with or without red light illumination (660 nm, 1 mW cm^-2^). SEAP production was profiled at 48 hours after illumination. Black block, with treatment; white block, without treatment. **e**, REDMAP_Cre_-catalyzed DNA recombination for fluorescent (EGFP) reporter expression. 6×10^4^ HEK-293T cells were co-transfected with pYZ208, pYZ247, and the Cre-dependent EGFP reporter (pDL78, P_hCMV_-*loxP*-STOP-*loxP*-EGFP-pA). Then, the transfected cells were supplied with PCB, followed by red light illumination (660 nm, 1 mW cm^-2^) for 48 hours. EGFP levels were profiled by fluorescence microscopy after illumination (using Image J for quantification). Representative images from 5 biological replicates; scale bar, 200 μm. Data in **b**, **c**, and **e** are expressed as means ± SD. Student’s *t-*tests were used for comparison. *n* = 3 or 5 independent experiments. Data in **d** is expressed as the mean ± SD. One-way ANOVA was used for multiple comparisons. *n* = 3 independent experiments. NS, not significant.

We designed and generated a series of fusion combinations of split-Cre fragments with ΔPhyA/FHY1 **(Fig. 1b)** and assessed their DNA recombination activities. To this end, we constructed a floxed-STOP SEAP reporter by inserting a STOP cassette, flanked by *loxP* sites, into the 5′ untranslated region (UTR) of *SEAP*; this design prevents reporter expression until the cassette is excised by Cre recombinase. We then used this floxed-STOP SEAP reporter to test the red light-inducible DNA recombination efficiency of all designed configurations of REDMAP_Cre_ in HEK-293T cells. The FHY1-CreN/ΔPhyA-CreC combination exhibited the highest fold increase (41-fold) of SEAP expression **(Fig. 1b)**. Notably, although this configuration demonstrated the most significant fold induction of Cre-*loxP* recombination, the absolute efficiency of Cre-*loxP* recombination upon irradiation remained low.

Using the FHY1-CreN/ΔPhyA-CreC configuration, we subsequently designed various linkers to connect FHY1 with CreN and ΔPhyA with CreC, aiming to achieve highly efficient DNA recombination. The combination of FHY1-L3-CreN and ΔPhyA-L5-CreC, hereafter termed REDMAP_Cre_ system, achieved the highest fold induction of 85-fold in SEAP expression upon red light illumination and PCB supplementation **(Fig. 1c** and **Supplementary Fig. 1)**. In the absence of either or both PCB and red-light illumination, no significant increase in SEAP expression was observed, confirming the requirement of both red light and PCB **(Fig. 1d)**. In addition, when HEK-293T cells were co-transfected with REDMAP_Cre_ and reporters for the enhanced green fluorescent protein (EGFP) **(Fig. 1e)** and luciferase **(Supplementary Fig. 2)**, both flanked by *loxP* sites, we observed a significant increase in the expression levels of either reporter upon red light illumination compared to conditions of darkness. Thus, we have successfully developed REDMAP_Cre_, enabling highly inducible DNA recombination in response to red light illumination and PCB supplementation.

### Characterization of DNA recombination control using REDMAP_Cre_

We subsequently introduced REDMAP_Cre_ into five commonly used rodent and human cell lines. SEAP expression was illumination-dependent in all cells tested (**Fig. 2a**), suggesting the broad applicability of REDMAP_Cre_. We also exposed REDMAP_Cre_ to different wavelengths of light and observed the highest extent of SEAP induction under 660 nm red light illumination, validating its specific responsiveness to red light (**Supplementary Fig. 3**). This system efficiently catalyzed the recombination of several mutant *loxP* sequences, as well as the original (non-mutant) *loxP* sequences, upon red light illumination (**Supplementary Fig. 4a-b**). Further, we evaluated REDMAP_Cre_’s capacity to excise STOP sequences of varying lengths (including 2 or 3 polyA sequences) flanked by *loxP* sequences, and to recombine a Cre-dependent <u>d</u>ouble-floxed inverted <u>o</u>pen reading frame (DIO) to express a SEAP reporter (**Supplementary Fig. 4c**). Each Cre-dependent reporter construct significantly increased SEAP expression following REDMAP_Cre_ induction, with the DIO reporter exhibiting the lowest basal recombination activity (**Supplementary Fig. 4d**).

**Figure 2.**
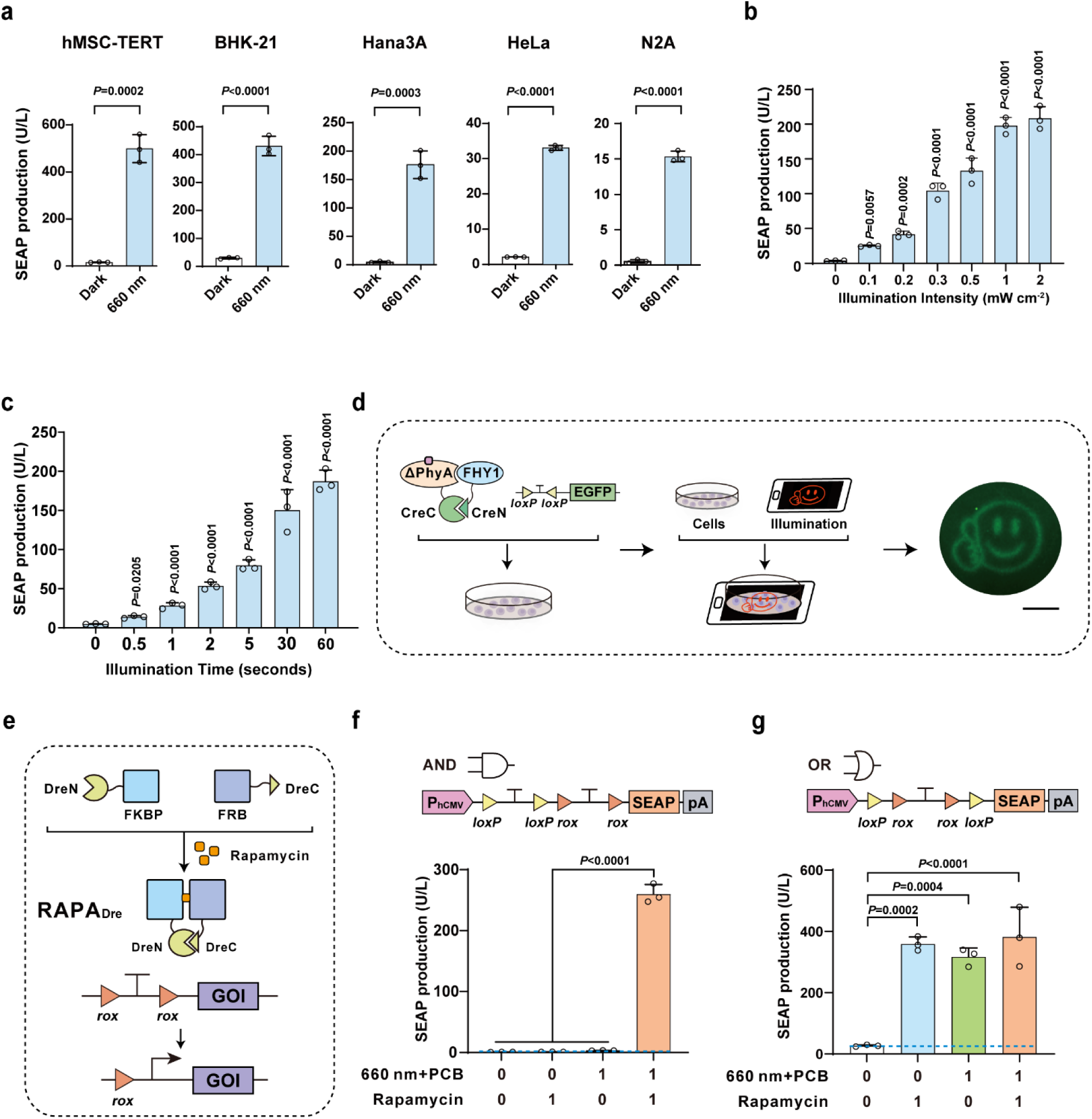
| Characterization of DNA recombination control with REDMAP_Cre_. **a**, REDMAP_Cre_-mediated SEAP expression in the indicated mammalian cell lines. Five mammalian cell lines (6×10^4^) co-transfected with pYZ208, pYZ247, and pGY125 were supplied with PCB and then illuminated with red light (660 nm, 1 mW cm^-2^) for 48 hours. SEAP production was quantified after illumination. **b**, Assessment of illumination-intensity-dependent REDMAP_Cre_ activity. HEK-293T cells (6×10^4^) transfected as described in **a** were supplied with PCB and then illuminated with red light (660 nm) at the indicated light intensities (0 to 2 mW cm^-2^) for one second. SEAP production was quantified 48 hours after illumination. **c**, Exposure-time-dependent REDMAP_Cre_ activity. HEK-293T cells (6×10^4^) transfected as described in **a** were supplied with PCB and then illuminated with red light (660 nm, 0.2 mW cm^-2^) for the indicated time (0 to 60 seconds). SEAP production was quantified 48 hours after illumination. **d**, Evaluation of the spatial control of REDMAP_Cre_-mediated recombination activity. HEK-293T cells (3×10^6^) were seeded into a 10-cm tissue culture dish and transfected with pYZ208, pYZ247, and pDL78. The dish was placed onto a smartphone screen projecting a pattern and illuminated with red light for 5 minutes (display brightness set to 100%, 10 μW cm^-2^) (schematic, left and middle). Fluorescence micrographs assessing EGFP production were taken 48 hours after illumination using fluorescence microscopy (bottom). Scale bar, 2 cm. **e**, Schematic representation of the rapamycin inducible Dre system (RAPA_Dre_). In the absence of rapamycin, Dre recombinase is divided into two inactive parts. In the presence of rapamycin, split Dre (1-246 aa/247-343 aa) reassociates to form a complete Dre based on rapamycin-dependent dimerization of FKBP and FRB, enabling excision of DNA sequences (STOP) flanked by *rox* sites. **f**, REDMAP_Cre_ and RAPA_Dre_ mediated SEAP expression (AND gate). The reporter was constructed with a *loxP* flanked STOP sequence followed by a *rox* flanked STOP sequence positioned upstream of *SEAP*, and SEAP should only be expressed when both STOP cassettes were successfully excised. HEK-293T cells (6×10^4^) transfected with plasmids encoding REDMAP_Cre_ and RAPA_Dre_ were induced by PCB plus red light or rapamycin as indicated. SEAP production was quantified 48 hours after induction. **g**, REDMAP_Cre_ and RAPA_Dre_ mediated SEAP expression (OR gate). The reporter was constructed with a STOP sequence flanked by *loxP* and *rox* sequences upstream of *SEAP*, and SEAP should be expressed when STOP cassettes were successfully excised. HEK-293T cells (6×10^4^) transfected with plasmids encoding REDMAP_Cre_ and RAPA_Dre_ system were induced by PCB plus red light or rapamycin as indicated. SEAP production was quantified 48 hours after induction. Data in **a**, **f**, and **g** are expressed as means ± SD. Student’s *t*-tests were used for comparison. *n* = 3 independent experiments. Data in **b, c** is expressed as the mean ± SD. One-way ANOVA was used for multiple comparisons. *n* = 3 independent experiments.

Next, we characterized the performance of the DNA recombination of REDMAP_Cre_ by transiently co-transfecting HEK-293T cells with vectors encoding REDMAP_Cre_ and a floxed-STOP SEAP reporter. We observed that SEAP production depended on the intensity and duration of illumination **(Fig. 2b-c)**. These results show the system’s high sensitivity to light: just 1 second of illumination at 0.3 mW/cm^2^ triggered approximately 50% of maximum DNA recombination, while peak SEAP expression was induced with only 1 second at 1 mW/cm^2^ **(Fig. 2b)**. We also compared the DNA recombination performance of REDMAP_Cre_ with those of two other available red/far-red light-inducible Cre recombinase systems, FISC^31^ and RedPA-Cre^32^. The DNA recombination efficiency of REDMAP_Cre_, indicated by the SEAP fold increase, was significantly higher than the other two when subjected to the same illumination intensity and duration, specifically 1 mW cm^−2^ for 1 second (**Supplementary Fig. 5**). Finally, we assessed the spatial control of DNA recombination activity of REDMAP_Cre_ by using photomask-mediated local illumination, revealing a distinct spatial pattern of DNA recombination, with evident EGFP activity in the regions only exposed to light **(Fig. 2d)**.

We then evaluated the compatibility of REDMAP_Cre_ with existing inducible recombinase systems by engineering recombinase-programmable AND and OR logic gates. Our initial step involved constructing a rapamycin-regulated Dre recombinase system, termed RAPA_Dre_, in which rapamycin induces the dimerization of FK506 binding protein (FKBP) and the FKBP–rapamycin binding (FRB) domain of the mammalian target of rapamycin (mTOR), thus reconstituting an active Dre recombinase capable of excising DNA sequences flanked by two *rox* sites **(Fig. 2e)**. To construct an AND logic gate, we inserted two STOP cassettes, each flanked by either *loxP* or *rox* sites (forming a *loxP*-STOP-*loxP*-*rox*-STOP-*rox* configuration), between the constitutive P_hCMV_ promoter and the SEAP reporter gene. According to this design, SEAP expression should occur only when both STOP cassettes are excised by the inducible Cre-*loxP* and Dre-*rox* recombinase systems. Our experiments demonstrated that SEAP production levels increased solely in the simultaneous presence of both inducers (660 nm light plus PCB and rapamycin) (**Fig. 2f**). We then developed an OR logic gate by introducing a STOP cassette, flanked by both *lox*P and *rox* sites (*loxP*-*rox*-STOP-*rox*-*loxP*), between the constitutive P_hCMV_ promoter and the SEAP reporter gene. In this OR gate configuration, the STOP cassette can be excised by either the Cre-*loxP* or the Dre-*rox* inducible recombinase system, resulting in SEAP expression under the control of the P_hCMV_ promoter. In the absence of both inducers (660 nm light plus PCB and rapamycin), SEAP leakage was minimal, while any combination of these inducers significantly enhanced SEAP expression (**Fig. 2g**). Collectively, these findings confirm that REDMAP_Cre_ is highly responsive to light, capable of rapid DNA recombination, and shows excellent compatibility with other inducible recombinase systems.

### REDMAP_Cre_-mediated DNA recombination in mice

We next assessed REDMAP_Cre_’s ability to induce DNA recombination *in vivo* on red light exposure. Vectors encoding REDMAP_Cre_ and a luciferase reporter were introduced into the livers of wild-type C57BL/6J mice via hydrodynamic tail vein (HTV) injection. Eight hours post-injection, mice were exposed to red light (660 nm, 20 mW cm^−2^) for one hour or not, with or without intraperitoneal injection of PCB **(Fig. 3a)**. Reporter expression was measured using an *in vivo* imaging system (IVIS). Mice that received both red light and PCB showed a marked increase in luciferase activity compared to controls exposed to only light, only PCB, or neither (**Fig. 3b**). Further experiments showed that even a single second of red-light exposure was sufficient to induce significant DNA recombination in mice (**Fig. 3c**).

**Figure 3.**
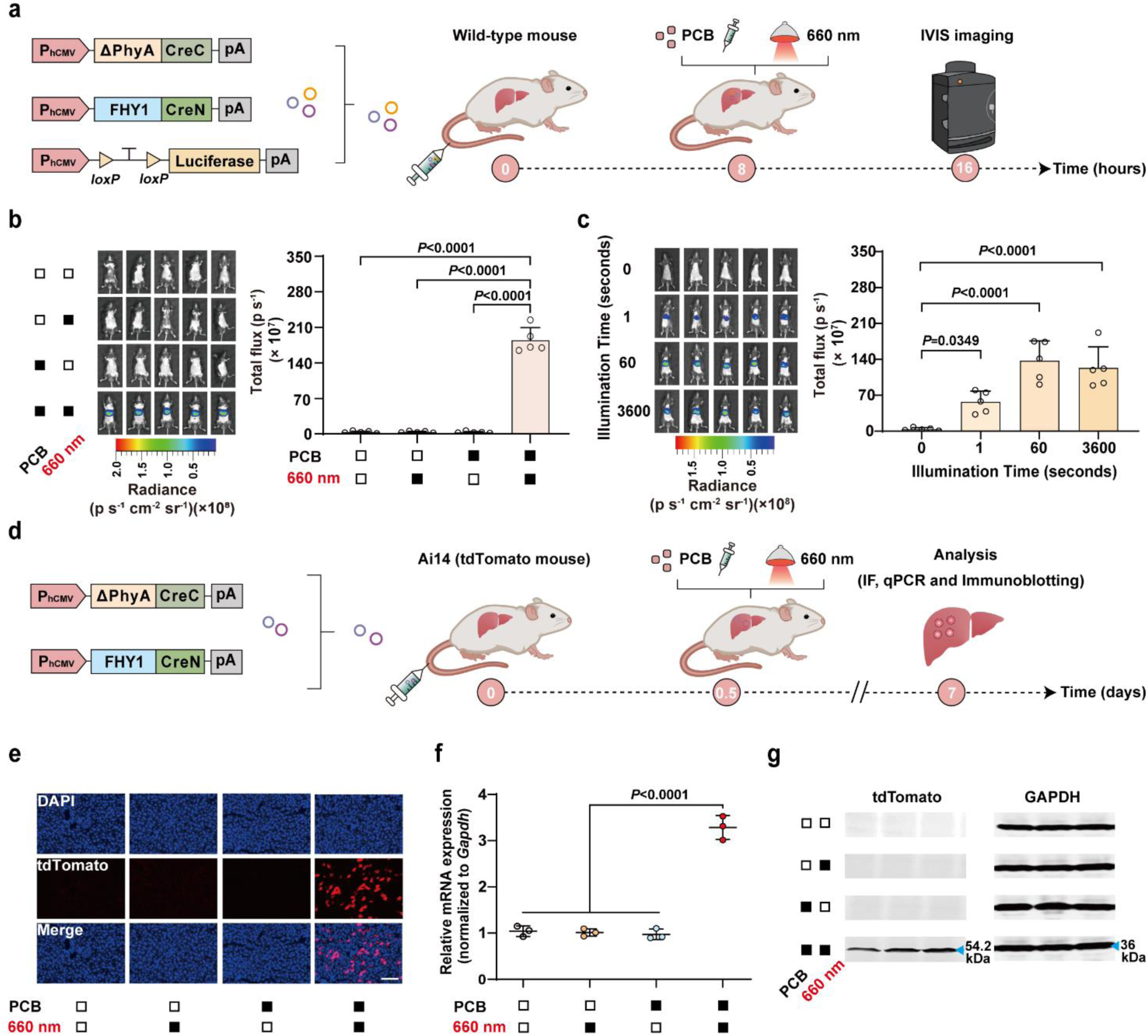
| DNA recombination with REDMAP_Cre_ in mice. **a**, Schematic representation of the genetic configurations of REDMAP_Cre_ (left) and the experimental procedure for REDMAP_Cre_-mediated gene recombination in mouse livers (right). **b**, REDMAP_Cre_-mediated gene recombination in wild-type C57BL/6J mouse livers. Mice were hydrodynamically injected (tail vein) with REDMAP_Cre_-encoding plasmids with Cre-dependent Luciferase reporter (pXY185, P_hCMV_-*loxP*-STOP-*loxP*-Luciferase-pA). At 8 hours post-injection, the mice were intraperitoneally injected with PCB (5 mg kg^-1^) and illuminated with red light (660 nm, 20 mW cm^-2^) for one hour. The control mice were exposed to either red light or PCB alone (or neither). Bioluminescence was quantified 8 hours after illumination using an *in vivo* imaging system (IVIS). **c**, Exposure-time-dependent REDMAP_Cre_ activity in mouse livers. Mice were hydrodynamically injected (tail vein) as described in **b**. At 8 hours post-injection, the mice were intraperitoneally injected with Bioluminescence was quantified 8 hours after illumination using IVIS. **d**, Schematic representation of the experimental procedure for REDMAP_Cre_-mediated gene recombination in *R26-tdTomato* transgenic (Ai14, *Gt(ROSA)26Sor^tm^*^14^*^(CAG-tdTomato)Hze^*) mouse livers. **e-g**, REDMAP_Cre_-mediated gene recombination in Ai14 tdTomato reporter mouse livers. Mice were hydrodynamically injected (tail vein) with the REDMAP_Cre_-encoding plasmids. At 12 hours post-injection, the mice were intraperitoneally injected with PCB (5 mg kg^-1^) and illuminated with red light (660 nm, 20 mW cm^-2^) for one second. The control mice were exposed to either red light or PCB alone (or neither). Mice were sacrificed, and livers were analyzed seven days after illumination. **e**, Representative fluorescence images of liver sections from the indicated groups. Blue, DAPI. Red, tdTomato. Scale bar, 100 μm. **f-g**, qPCR (**f**), and immunoblotting (**g**) analysis of tdTomato in isolated liver tissues. Black block, with treatment; white block, without treatment. Data in **b-c, f-g** are expressed as means ± SEM. One-way ANOVA was used for multiple comparisons. *n* = 3-5 mice.

We subsequently conducted tests on the induction of DNA recombination at a specific genomic locus by REDMAP_Cre_. Plasmids encoding REDMAP_Cre_ were delivered into adult *R26-tdTomato* transgenic mice (Ai14, *Gt(ROSA)26Sor^tm^*^14^*^(CAG-tdTomato)Hze^*)^34^ via HTV injection. These mice harbored a Cre reporter allele, which included a *loxP*-flanked STOP cassette. This cassette was designed to prevent the transcription of a CAG promoter-driven tdTomato reporter until excised by Cre recombinase. Twelve hours after injection, the Ai14 tdTomato reporter mice were intraperitoneally injected with PCB (0 or 5 mg kg^-1^) and exposed to red light (660 nm, 20 mW cm^-2^) for one second or not **(Fig. 3d)**. One week after illumination, fluorescence imaging was used to measure the tdTomato signals from isolated mouse livers. This revealed a significant induction of the tdTomato signal exclusively in mice that were both injected with PCB and exposed to red light **(Fig. 3e)**. qPCR and immunoblotting analyses of isolated liver tissues revealed that mice subjected to both red light illumination and PCB supplementation exhibited significantly stronger tdTomato signals compared to control mice subjected to red light, PCB alone, or neither (**Fig. 3f-g)**. These findings demonstrate that REDMAP_Cre_ can activate DNA recombination in mice with only one second of red-light exposure.

### AAV-REDMAP_Cre_-mediated DNA recombination in mouse liver and muscle

We employed AAV-mediated delivery for DNA recombination in mice, utilizing different serotypes to target specific tissues *in vivo*^35^. Specifically, we incorporated our REDMAP_Cre_ system into the liver tropism serotype AAV8, to enable the constitutive expression of its components, FHY1-CreN and ΔPhyA-CreC. These vectors were subsequently transduced into the livers of the adult Ai14 tdTomato reporter mice via tail vein injection. Two weeks following the AAV injection, the transgenic mice were exposed to red light (660 nm, 20 mW cm^-2^) for one hour, coupled with an intraperitoneal injection of PCB **(Supplementary Fig. 6a)**. One week after this treatment, the mice were euthanized, and their livers were harvested for the measure of the tdTomato signals. Fluorescence microscopy imaging of these sections showed that Cre-*loxp* recombination-mediated tdTomato expression occurred exclusively in mice exposed to both PCB and red light. In contrast, no tdTomato signal was detected in mice that were not treated with PCB or exposed to light **(Supplementary Fig. 6b)**. The qPCR and immunoblotting analysis also indicated that Cre-*loxp* recombination-mediated tdTomato expression occurred exclusively in mice subjected to both PCB and light exposure **(Supplementary Fig. 6c-d)**.

We subsequently assessed the AAV-REDMAP_Cre_-mediated DNA recombination in the muscles of mice. For this purpose, we chose AAV9, known for its muscle tropism, as the serotype for the backbone vector to express the REDMAP_Cre_ components. These AAV-REDMAP_Cre_ viral vectors were injected into the left gastrocnemius muscles of adult Ai14 tdTomato reporter mice via intramuscular injection. Two weeks after the AAV injection, the transgenic mice received an intraperitoneal injection of PCB, followed by exposure to red light (660 nm, 20 mW cm^-2^) for one hour **(Fig. 4a)**. One week after light exposure, the mice were euthanized, and their left gastrocnemius muscles were harvested for the measure of the tdTomato signals. Fluorescence microscopy, qPCR, and immunoblotting analysis indicated that Cre-*loxp* recombination-mediated tdTomato expression occurred exclusively in mice subjected to both PCB and light exposure **(Fig. 4b-d)**.

**Figure 4.**
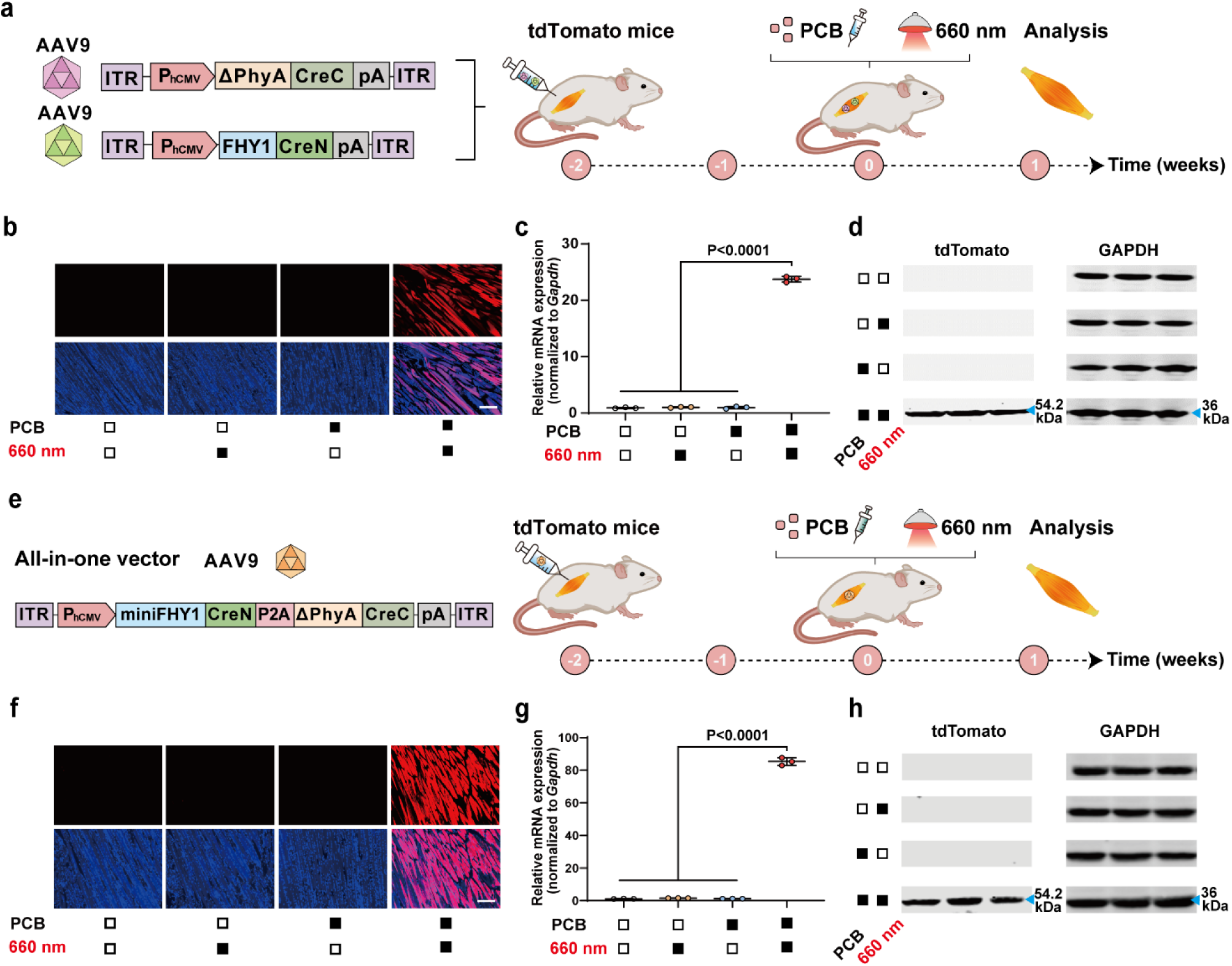
| AAV-REDMAP_Cre_-mediated DNA recombination in mouse muscles. **a**, Schematic representation of the experimental procedure for AAV9 delivery of REDMAP_Cre_ in Ai14 tdTomato reporter mouse left gastrocnemius muscles. Mice were intramuscularly injected with a mixture of AAV encoding REDMAP_Cre_ [(pYZ590 (ITR-P_hCMV_-ΔPhyA-CreC-pA-ITR) and pYZ591 (ITR-P_hCMV_-FHY1-CreN-pA-ITR)]. After two weeks, mice were intraperitoneally injected with PCB (20 mg kg^-1^) and illuminated with red light (660 nm, 20 mW cm^-2^) for one hour. The mice were sacrificed, and the muscles were analyzed seven days after illumination. **b**, Representative fluorescence images of muscle sections from the indicated groups. Blue, DAPI. Red, tdTomato. Scale bar, 200 μm. **c-d**, qPCR (**c**), and immunoblotting (**d**) analysis of tdTomato in isolated muscle tissues. **e**, Schematic depicting the genetic configuration of the all-in-one AAV vector constructs and the experimental procedure for delivery in Ai14 tdTomato reporter mouse left gastrocnemius muscles. Mice were intramuscularly injected with an all-in-one AAV encoding REDMAP_Cre_ [pYZ751 (ITR-P_hCMV_-miniFHY1-CreN-P2A-ΔPhyA-CreC-pA-ITR)]. Two weeks after AAV injection, the transgenic mice were exposed to both PCB and red light, and the gastrocnemius muscles were isolated and groups. Blue, DAPI. Red, tdTomato. Scale bar, 200 μm. **g-h**, qPCR (**g**) and immunoblotting (**h**) analysis of tdTomato in isolated muscle tissues. Black block, with treatment; white block, without treatment. Data in **c** and **g** are expressed as means ± SEM. One-way ANOVA was used for multiple comparisons. *n* = 3 mice.

The adoption of a single AAV vector has emerged as a promising delivery strategy for enhancing transduction efficiency in *in vivo* applications^36,37^. To overcome the inherent packaging limitations of a single AAV vector, we meticulously refined the REDMAP_Cre_ construct by removing the N-terminal amino acids of FHY1. Notably, our experimental results demonstrated that truncating the first 38 amino acids of FHY1 (yielding miniFHY1) maintains the robust DNA recombination activity of REDMAP_Cre_ (**Supplementary Fig. 7**). Subsequently, we designed an all-in-one AAV vector, encoding the REDMAP_Cre_ components linked by the self-cleaving P2A peptide (ITR-P_hCMV_-miniFHY1-CreN-P2A-ΔPhyA-CreC-pA-ITR). This vector was injected into the left gastrocnemius muscles of adult Ai14 tdTomato reporter mice. Two weeks after injection, the transgenic mice were exposed to PCB and red light **(Fig. 4e)**. One week following the treatment, the left gastrocnemius muscles were isolated for analysis. Fluorescence microscopy, as well as qPCR and immunoblotting analysis, showed mice exposed to both PCB and red light exhibited a significant increase in the tdTomato signal compared to other control groups **(Fig. 4f-h)**. This all-in-one AAV vector was also delivered into mouse livers and demonstrated robust efficiency of REDMAP_Cre_ mediated DNA recombination (**Supplementary Fig. 8**). Collectively, these findings demonstrate that REDMAP_Cre_, when delivered by AAV vectors, results in significant DNA recombination in the livers and muscles of mice exposed to both PCB and red light.

### Generation and characterization of the REDMAP_Cre_ mouse line

To facilitate studies on the functional implications of gene manipulation *in vivo*, we generated a transgenic mouse line by inserting the REDMAP_Cre_ gene cassette into the *Rosa26* locus using CRISPR-Cas9^38^, enabling generalized expression across various tissues **(Fig. 5a)**. Successful insertion was confirmed through PCR amplification of both the left and right boundaries between *Rosa26* and the REDMAP_Cre_ gene cassette, followed by DNA sequencing (**Fig. 5b-c**). To assess the functionality of the REDMAP_Cre_ transgenic mouse line, we transduced these mice with an AAV8 virus carrying the *loxP*-STOP-*loxP*-Luciferase reporter via tail-vein injection. Two weeks after AAV injection, the mice were exposed to red light (660 nm, 20 mW cm^-2^) for one hour or with no exposure, with or without an intraperitoneal injection of PCB (20 mg kg^-1^). Luciferase reporter expression was measured using IVIS one week after illumination **(Fig. 5d)**. Only the mice exposed to both PCB and red light exhibited a significant increase in luciferase signal compared to other control mice **(Fig. 5e-f)**. The data indicate that the REDMAP_Cre_ transgenic mouse line was successfully generated.

**Figure 5.**
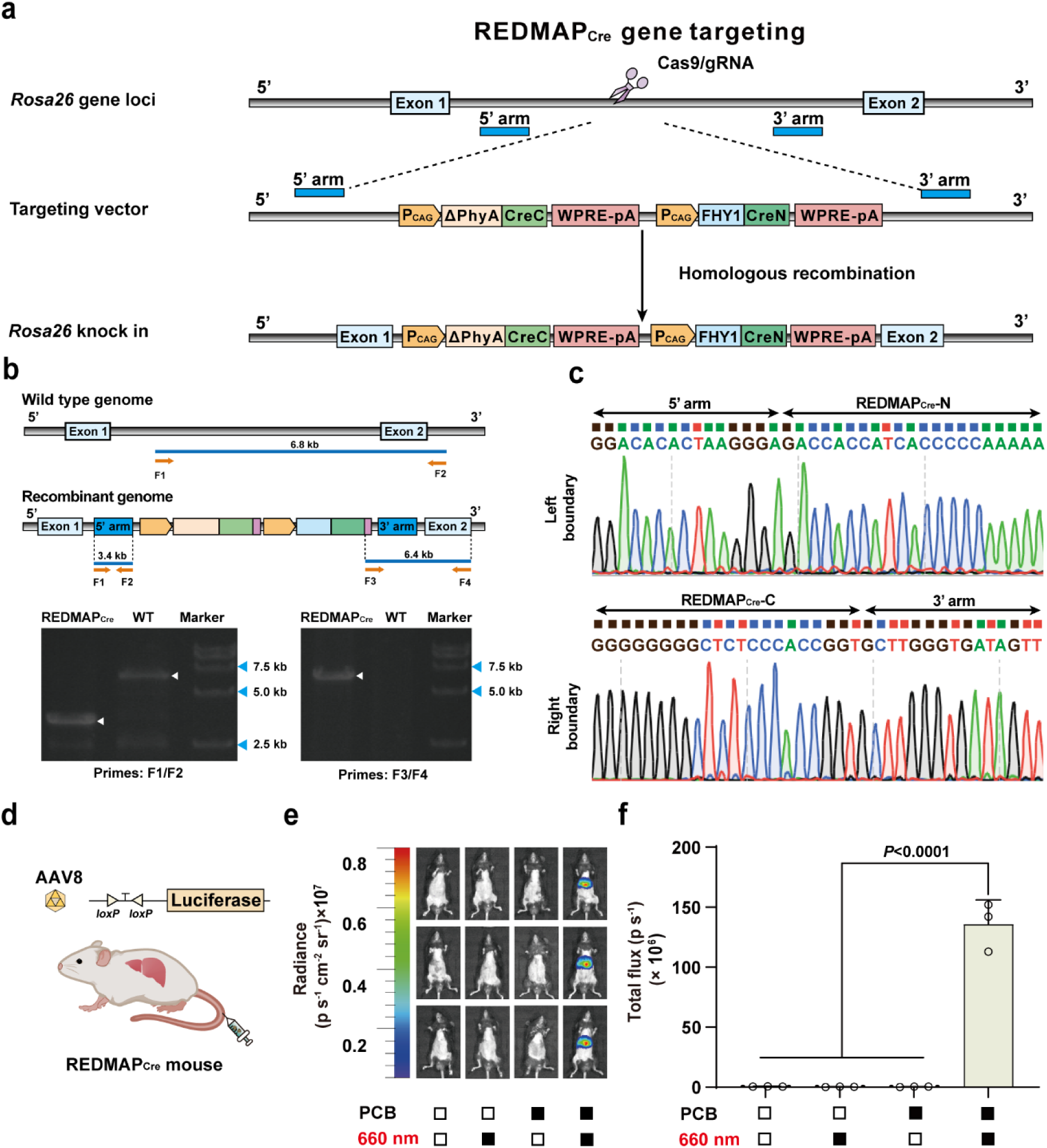
| Generation and characterization of REDMAP_Cre_ mice. **a**, Schematic for the REDMAP_Cre_ mice targeting strategy. Using CRISPR/Cas9, the expression frame of P_CAG_-ΔPhyA-CreC-WPRE-pA-P_CAG_-FHY1-CreN-WPRE-pA was inserted into the *Rosa26* gene locus through homologous recombination. **b**, Identification of REDMAP_Cre_ mice by PCR (top) and the representative electrophoresis images to validate REDMAP_Cre_ insertion (bottom). The primers (F1, F2) were designed to identify the positive genome in the 5’ arm, and the primers (F3, F4) were designed to identify the positive genome in the 3’ arm. **c**, Sequences of boundaries between *Rosa26* and the REDMAP_Cre_cassette. **d**, Schematic representation of AAV8 delivery of a Cre-dependent reporter in REDMAP_Cre_ mice. **e-f**, AAV8 (ITR-P_hCMV_-*loxP*-STOP-*loxP*-Luciferase-pA-ITR) were transduced into the REDMAP_Cre_ mice through tail vein injection. Two weeks after AAV injection, the mice were intraperitoneally injected with PCB (20 mg kg^-1^) and then illuminated with red light (660 nm, 20 mW cm^-2^) for one hour. The control mice were exposed to either red light or PCB alone (or neither). Bioluminescence was quantified one week after illumination using IVIS. Black block, with treatment; white block, without treatment. Data in **f** are expressed as means ±SEM. One-way ANOVA was used for multiple comparisons. *n* = 3 mice.

To further test whether the REDMAP_Cre_ mouse line enables the induction of DNA recombination in response to red light, REDMAP_Cre_ transgenic mice were crossed with Ai14 tdTomato reporter mice. The latter carry the *loxP*-STOP-*loxP*-tdTomato reporter sequences, allowing us to obtain double transgenic mouse line REDMAP_Cre_:Ai14 **(Fig. 6a)**. To assess *in vitro* DNA recombination, we isolated fibroblasts, bone marrow-derived stem cells (BMSCs), and cortical cells from REDMAP_Cre_:Ai14 double transgenic mice, and exposed these cells to red light in the presence of PCB (**Supplementary Fig. 9a-b**). Fluorescence imaging revealed that only cells exposed to both PCB and red light exhibited an increased tdTomato signal; in contrast, no significant reporter induction was observed in other control conditions (**Supplementary Fig. 9c-e**). Moreover, we observed that Cre recombinase activity depended on illumination intensity (**Fig. 6b**) and exposure time (**Fig. 6c**). A mere one second of red-light stimulation at 0.2 mW cm^−2^ was sufficient to drive tdTomato expression (∼423-fold) *in vitro* (**Fig. 6c**). Furthermore, we observed a plateau of induced tdTomato expression reaching an approximate 1578-fold increase, following a 5-second exposure to an illumination intensity of 1 mW cm^-2^ **(Fig. 6b)**.

**Figure 6.**
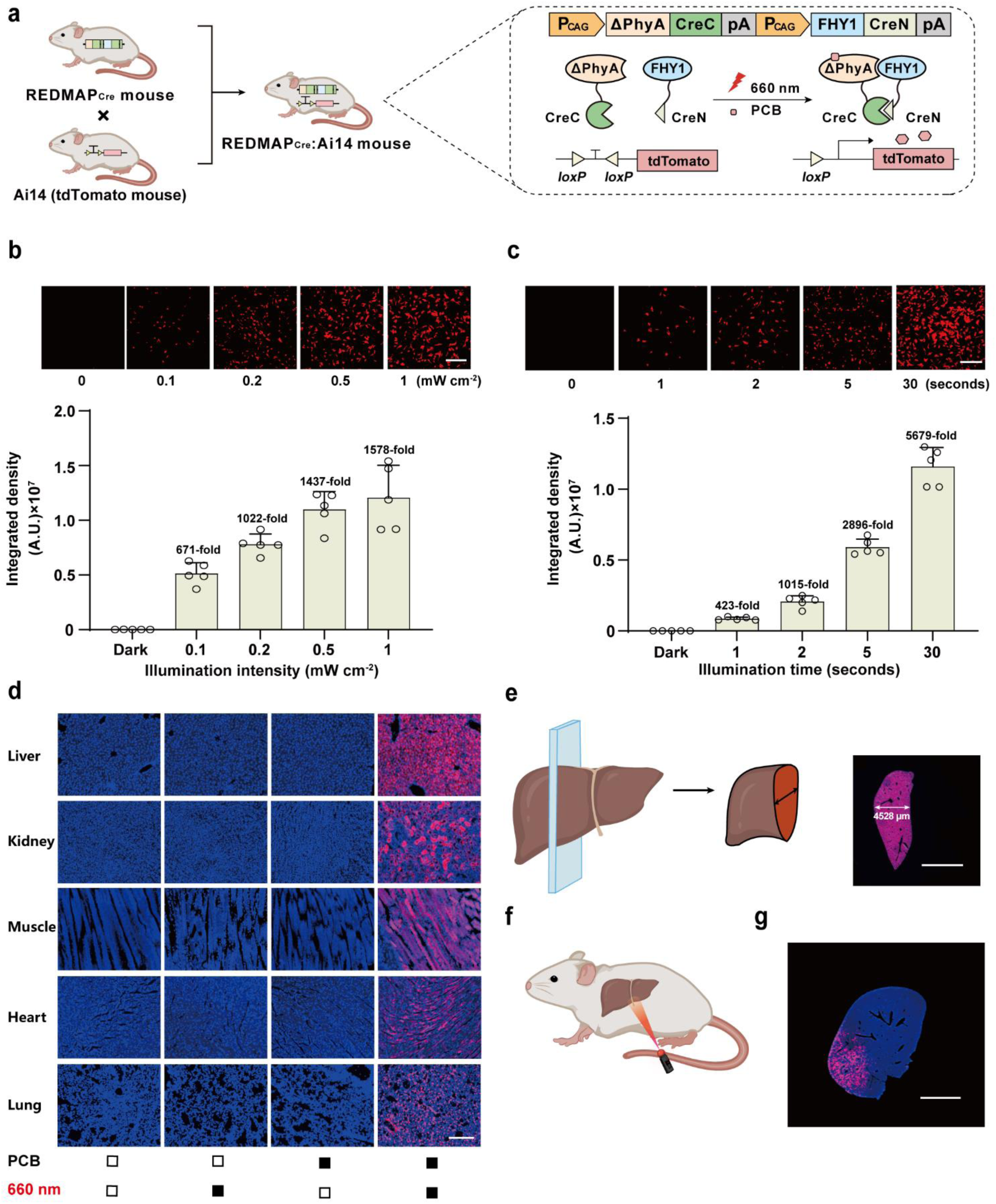
| Generation of REDMAP_Cre_:Ai14 transgenic mice and characterization of gene recombinase activity. **a**, Schematic for red-light dependent activation of the tdTomato reporter in REDMAP_Cre_:Ai14 mice. The transcriptional stop cassette flanked by two *loxP* sites was excised by REDMAP_Cre_ upon PCB injection and red-light illumination. The following experiments were conducted in double-heterozygous REDMAP_Cre_:Ai14 mice. **b**, Exposure intensity-dependent REDMAP_Cre_ activity. Fibroblasts isolated from mouse skin were illuminated with red light (660 nm) for 5 seconds at the indicated intensity. Fluorescent tdTomato images were acquired by fluorescence microscopy 48 hours after illumination (using Image J for quantification). Red, tdTomato. Scale bar, 500 μm. The graph at the bottom represents the quantification of tdTomato fluorescence intensity under the indicated conditions. **c**, Exposure time-dependent REDMAP_Cre_ activity. Fibroblasts isolated from mouse skin were illuminated with red light (660 nm, 0.2 mW cm^-2^) for the indicated times. Red, tdTomato. Scale bar, 500 μm. Data in **b-c** are expressed as means ± SD. *n* = 5 independent experiments. **d**, Representative fluorescence images of various tissue sections from REDMAP_Cre_:Ai14 mice. Adult mice were intraperitoneally injected with PCB (200 mg kg^-1^) and illuminated with red light (660 nm, 20 mW cm^-2^) for one hour. The mice were sacrificed, and the tissues were harvested and analyzed seven days after illumination. Black block, with treatment; white block, without treatment. Blue, DAPI. Red, tdTomato. Scale bar, 200 μm. **e**, Tissue penetration of REDMAP_Cre_:Ai14 mice in livers. Mice were treated as described in **d**, and livers were harvested seven days after illumination and sectioned as indicated (left). Representative fluorescence images of liver sections were obtained using a Digital Pathology System (right). Blue, DAPI. Red, tdTomato. Scale bar, 5 mm. **f**, Spatial control gene recombination in REDMAP_Cre_:Ai14 mouse livers. The mice were locally irradiated using a laser pen, and the livers were harvested and analyzed seven days after illumination. **g**, Representative fluorescence image of liver sections obtained using a Digital Pathology System. Blue, DAPI. Red, tdTomato. Scale bar, 5 mm.

To evaluate the *in vivo* DNA recombination activity across various tissues such as the heart, liver, and spleen, among others, in the double transgenic mouse line REDMAP_Cre_:Ai14, six-week-old REDMAP_Cre_:Ai14 mice were intraperitoneally injected with PCB followed by exposure to red light (660 nm, 20 mW cm^-2^). Subsequently, the harvested tissues were dissected and sectioned for fluorescence imaging. A tdTomato signal was observed in all examined internal tissues, including the liver, kidney, muscle, heart, lung, brain, intestine, spleen, and also in the skin of the mice. No background activity was detected in tissues in the absence of red light or PCB **(Fig. 6d** and **Supplementary Fig. 10)**. Notably, fluorescent reporter signals were observed throughout longitudinal liver sections, indicating widespread DNA recombination in the liver (**Fig. 6e**). We further performed localized irradiation of the liver from the mouse abdomen using a 660 nm laser pen (**Fig. 6f**). Subsequently, the liver sections exhibited localized increases in tdTomato fluorescent signals, confirming that DNA recombination occurred exclusively in areas received red light illumination (**Fig. 6g**). Taken together, these findings underscore that the REDMAP_Cre_ mouse line enables highly efficient and spatially targeted gene recombination across various tissues *in vivo*.

### Optogenetic induction of metabolic dysregulation and cell ablation in mice

Having demonstrated the deep tissue penetration capability REDMAP_Cre_, we next applied this system to investigate gene function in deep tissues. The ubiquitin-like with PHD and ring finger domains 1 (UHRF1) has been shown to promote fatty acid and protein synthesis by suppressing AMP-activated protein kinase (AMPK)^39^. To examine the role of UHRF1 in mouse liver, REDMAP_Cre_ mice were crossed with Rosa-LSL-UHRF1 mice, in which Cre-induced recombination drives *UHRF1* expression (**Fig. 7a)**. After confirming successful recombination through genotyping, the resulting REDMAP_Cre_:Rosa-LSL-UHRF1 offspring were used for further functional analyses. Five-week-old REDMAP_Cre_:Rosa-LSL-UHRF1 mice were intraperitoneally injected with PCB and subsequently exposed to red light (660 nm, 20 mW cm^-2^) for one hour. One week after illumination, these mice exhibited significantly higher fed and fasted blood glucose levels compared to controls (**Fig. 7b-c**), as well as insulin resistance, as indicated by glucose and insulin tolerance tests (**Fig. 7d-e**). Moreover, PCB– and light-treated mice showed elevated liver triacylglycerol (TG) and serum TG levels (**Fig. 7f-g**) and increased hepatic lipid deposition (**Fig. 7h**), suggesting that optogenetic overexpression of UHRF1 induced insulin resistance and hepatic lipid accumulation.

**Figure 7.**
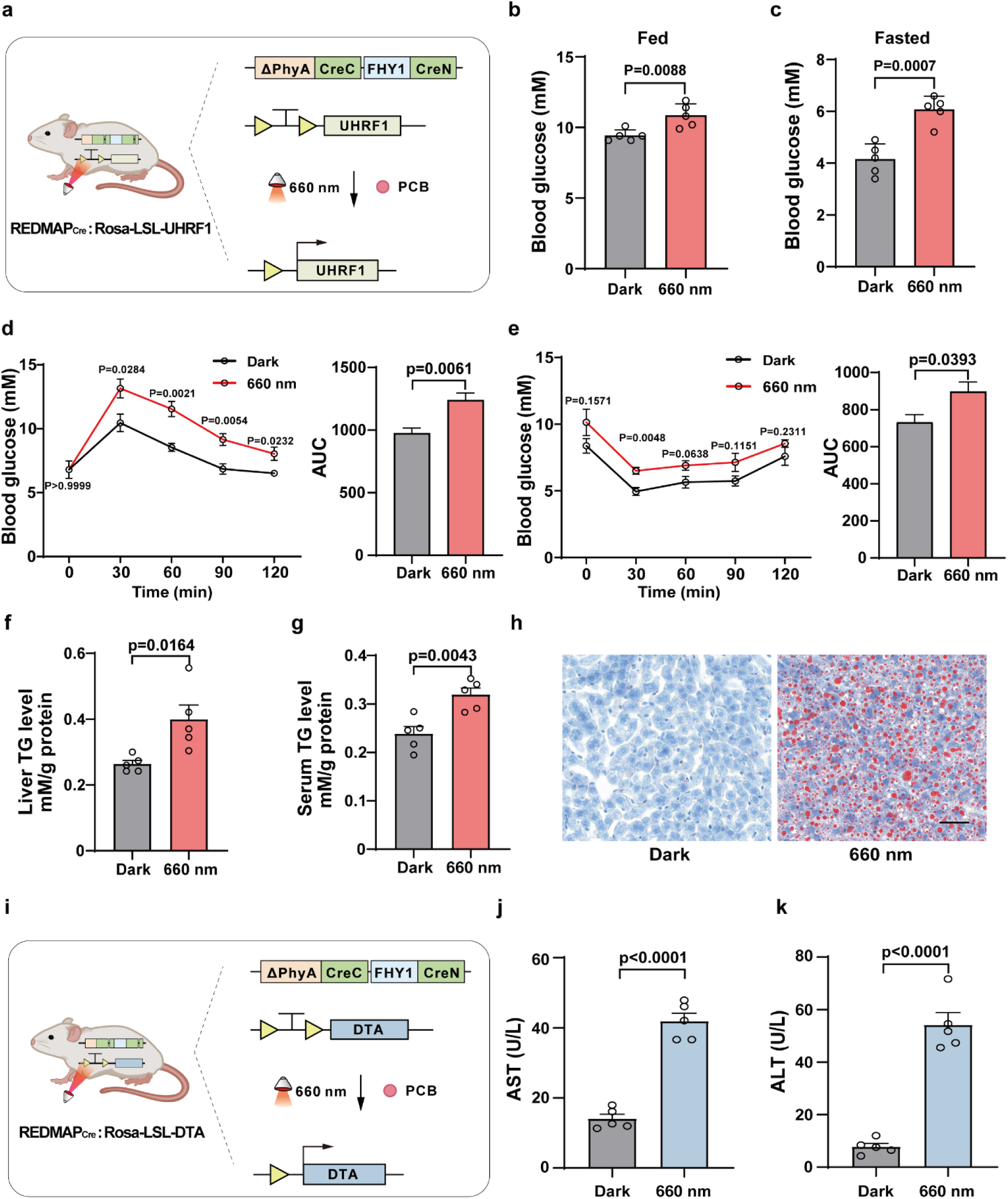
| Optogenetic induction of metabolic dysregulation and cell ablation in mice. **a**, Schematic for red-light-dependent activation of UHRF1 in REDMAP_Cre_:Rosa-LSL-UHRF1 mice. Upon PCB injection and red-light illumination, REDMAP_Cre_ excised the transcriptional stop cassette flanked by two *loxP* sites, enabling UHRF1 expression. **b-h**, REDMAP_Cre_:Rosa-LSL-UHRF1 mice were intraperitoneally injected with PCB (20 mg kg^-1^) and exposed to red light (660 nm, 20 mW cm^-2^) for one hour. Blood glucose levels were measured in the fed **(b)** and fasted (**c**) states on day 7 post-illumination. Glucose tolerance tests (**d**) and insulin tolerance tests (**e**) were conducted on day 8 and day 10, respectively. Mice were sacrificed for analysis of liver triglyceride (TG) (**f**) and serum TG (**g**) levels. (**h**) Liver samples were subjected to Oil Red O staining. Scale bar, 50 μm. **i**, Schematic for red-light-dependent activation of the diphtheria toxin fragment A (DTA) in REDMAP_Cre_:Rosa-LSL-DTA mice. **j-k**, REDMAP_Cre_:Rosa-LSL-DTA mice were intraperitoneally injected with PCB (20 mg kg^-1^) and illuminated with red light (660 nm, 20 mW cm^-2^) for one hour. Serum aspartate aminotransferase (AST) (**j**) and alanine aminotransferase (ALT) (**k**) levels were measured 24 hours post-illumination. Data in **b-g** and **j-k** are presented as means ± SEM. Statistical comparisons were performed using Student’s *t-*tests. *n* = 5 mice.

Targeted cell ablation is essential for understanding the functional roles of specific cell populations to tissue development, homeostasis, and regeneration^40^. However, achieving non-invasive, remote-controlled cell ablation in deep tissues remains a challenge. To future validate the capability of REDMAP_Cre_ for light-induced cell ablation, REDMAP_Cre_ mice were crossed with Rosa-LSL-DTA mice (**Fig. 7i**). The diphtheria toxin fragment A (DTA) subunit specifically inhibits protein synthesis, leading to apoptosis via activation of mitochondrial-dependent apoptotic pathways^41^. After confirming successful recombination through genotyping, the resulting REDMAP_Cre_:Rosa-LSL-DTA offspring were used for functional analyses. Six-week-old REDMAP_Cre_:Rosa-LSL-DTA mice were intraperitoneally injected with PCB and exposed to red light (660 nm, 20 mW cm^-2^) for one hour. Twenty-four hours post-illumination, serum levels of aspartate aminotransferase (AST) and alanine aminotransferase (ALT)—established biomarkers of hepatic injury—were significantly elevated in PCB– and light-treated mice compared to controls (**Fig. 7j-k**), confirming successful optogenetic induction of hepatocyte ablation. In summary, these findings demonstrate that the REDMAP_Cre_ transgenic mouse model enables efficient, non-invasive, and spatiotemporally controlled gene regulation in vivo and serves as a valuable tool for studying gene function with high precision and control.

## Discussion

In this study, we present REDMAP_Cre_, a red light-controlled split-Cre recombinase system to overcome limitations of existing optogenetic recombinases. By engineering a ΔPhyA/FHY1 heterodimerization module responsive to 660 nm light^33^, REDMAP_Cre_ enables spatiotemporal control of genomic recombination, with its activity tunable by adjusting the intensity and duration of light exposure. The system exhibits rapid activation within seconds, high recombination efficiency, and negligible background activity in the absence of both phycocyanobilin (PCB) and illumination. Notably, the compact design of REDMAP_Cre_ (<4.7 kb) allows for single-AAV delivery, facilitating efficient in vivo applications with minimal vector burden. Furthermore, we established a REDMAP_Cre_ transgenic mouse model, demonstrating remote– and traceless-controlled DNA recombination in the liver and enabling optogenetic induction of insulin resistance and hepatic lipid deposition via Cre-dependent *UHRF1* overexpression and targeted cell ablation through DTA expression. This REDMAP_Cre_ system offers a powerful tool for dissecting the roles of specific cell populations in various physiological and pathological processes, with potential applications in regenerative medicine, disease modeling, and therapeutic interventions.

Our findings highlight REDMAP_Cre_ as a powerful tool for spatiotemporal gene recombination, enabling targeted DNA modification in specific cell types and tissues. Compared to conventional photoactivatable Cre models (e.g., PA-Cre 3.0, ePA-Cre, DiLi_Cre_) that rely on blue/UV light (low tissue penetration)^21,22,29,42–44^, REDMAP_Cre_ harnesses the superior tissue penetration of red light^25,45^ facilitating efficient recombination in deep-seated organs such as the liver. This expanded capability makes REDMAP_Cre_ particularly valuable for positionally restricted mosaic analyses and the generation of localized cancer models via optogenetic oncogene activation^46^. Moreover, REDMAP_Cre_ is compatible with other recombinases, enabling the development of dual-recombinase systems through crossbreeding with transgenic lines harboring orthogonal recombinases. This approach could significantly enhance in vivo lineage tracing, offering deeper insights into complex biological phenomena such as biliary epithelial cell fate transitions^47,48^ and cardiomyocyte regeneration following injury^49,50^.

A persistent challenge in photoactivatable Cre systems is unintended recombination due to ambient light exposure, particularly in external tissues like the skin in transgenic mice^22,28,29^. To mitigate this issue, researchers often impose stringent light-shielding measures, which disrupt natural light cycles and complicate studies in chronobiology. REDMAP_Cre_ addresses this limitation through a dual-input activation mechanism, requiring both PCB administration and red-light illumination for recombination. By decoupling activation from ambient light exposure, REDMAP_Cre_ eliminates the need for light deprivation protocols, allowing experimental animals to maintain physiological circadian rhythms under standard housing conditions.

Looking ahead, integrating additional site-specific recombinases such as Dre recombinase and Flp recombinase, as well as nucleases like CRISPER-associated protein 9 (Cas9) and CRISPER-associated protein 12 (Cas12a), into photoactivatable transgenic models could transform our ability to study cellular dynamics and lineage fate decisions. With its noninvasive, highly specific, and precisely controllable gene recombination capabilities, REDMAP_Cre_ holds immense potential for genome engineering applications in both basic research and biomedical sciences.

## Methods

### Cloning and plasmid construction

Some expression vectors are constructed using a MultiS One Step Cloning kit (C113-01, Vazyme) according to the manufacturer’s instructions. All relevant plasmids have been confirmed by sequencing (Shanghai Personalbio Technology).

### Cell culture and transfection

Human embryonic kidney 293T cells (HEK-293T, CRL-11268, ATCC), human cervical adenocarcinoma cells (HeLa, CCL-2, ATCC), HEK-293-derived Hana3A cells engineered for constitutive expression of RTP1, RTP2, REEP1, and Gαoλϕ, Baby hamster kidney cell line (BHK-21, CCL-10, ATCC), mouse Neuro-2a cells (N2A, CCL-131, ATCC) and telomerase-immortalized human mesenchymal stem cells (hMSC-TERT) were cultured in Dulbecco′s Modified Eagle Medium (DMEM; C11995500BT, Gibco) containing 10% (v/v) fetal bovine serum (FBS; FBSSA500-S, AusGeneX) and 1% (v/v) penicillin/streptomycin solution (ST488, Beyotime). All the cell lines were cultured at 37 °C in a humidified atmosphere containing 5% CO_2_ and have been regularly tested for the absence of *Mycoplasma* and bacterial contamination.

HEK-293T, HeLa, and Hana3A cells were transfected with an optimized polyethyleneimine (PEI)-based protocol. Briefly, the cells were plated 6×10^4^ cells per well in a 24-well cell culture plate and cultured for 18 hours before transfection. Subsequently, cells were incubated for six hours with 50 µl of PEI and DNA mixture at a mass ratio of 3:1 (PEI, molecular weight 40,000, stock solution 1 mg mL^−1^ in ddH_2_O; 24765, Polysciences). For the transfection of hMSC-TERT, BHK-21, and N2A, cells were plated into a 24-well plate and cultivated overnight to 70–80% confluency at the time of transfection with Lipo8000 Transfection Reagent (C0533, Beyotime) according to the manufacturer’s protocol.

### SEAP reporter assay

The quantification of human placental SEAP in a cell culture medium was conducted using a *p*-nitrophenylphosphate-based light absorbance time course^51^. Briefly, 120 µL of substrate solution was employed, consisting of 100 µL of 2×SEAP buffer containing 20 mM homoarginine, 1 mM MgCl_2_, 21% (v/v) diethanolamine at pH 9.8, and an additional 20 µL of substrate solution containing 120 mM p-nitrophenyl phosphate. This solution was added to 80 µL of cell culture supernatant that had been heat-inactivated (65 °C, 30 minutes). The absorbance time course at 405 nm was monitored at 37 °C by using a Synergy H1 hybrid multimode microplate reader (BioTek) equipped with Gen5 software (version 2.04). The quantification of SEAP production was calculated based on the slope of the time-dependent increase in light absorbance.

### Luciferase reporter assay

Luciferase activity levels were assayed using the Firefly Luciferase Reporter Gene Assay Kit (RG005, Beyotime) according to the manufacturer’s instructions. Briefly, the cells were harvested and treated with 100 μL cell lysate each well for 2 minutes. Then, the cells were centrifuged at 12,000 rpm for 5 minutes, and the supernatants were collected. The 20 μL luciferin was added to the 20 μL supernatant, and the signal was detected using the Synergy H1 hybrid multi-mode microplate reader (BioTek).

### Fluorescence imaging and fluorescence intensity measurements

Fluorescence images of EGFP or tdTomato expression cells were performed with an inverted fluorescence microscope (Olympus IX71, TH4-200, Olympus) equipped with an Olympus digital camera (Olympus DP71, Olympus), a 495/535-nm (B/G/R) excitation/emission filter set. Fluorescence intensity measurements were analyzed using Image J (version 1.8.0).

### Extraction and purification of phycocyanobilin

Phycocyanobilin (PCB) was obtained from Frontier Scientific (P14137) or extracted from phycocyanin according to a modified protocol^52^. Ten grams of phycocyanin powder (R3, Binmei Biotechnology) was boiled with 100 mL of methanol for 6 hours. The liquid was cleared by filtration, concentrated in a rotary evaporator to 20 mL, and mixed with 20 mL of chloroform and 40 mL of ddH_2_O. The PCB-containing organic phases were collected and evaporated in vacuo, dissolved in 200 μL of dimethyl sulfoxide (DMSO), and stored at –20 ℃. The PCB concentration was determined by diluting the DMSO stock 1:100 into 1 mL 95% MeOH and 5% hydrochloric acid (HCl, 37.5%). The absorbance was then measured at 680 nm by using a Synergy H1 hybrid multimode microplate reader (BioTek) equipped with Gen5 software (version 2.04)^53^. The PCB concentration was calculated based on the standard curve established by the known concentration of PCB.

### DNA recombination performance of REDMAP_Cre_ in mammalian cells

The 6×10^4^ HEK-293T cells were plated in a 24-well plate and transfected with plasmids encoding REDMAP_Cre_ and a Cre-dependent reporter. Twenty-four hours after the transfection, the cells were supplemented with 5 μM PCB in the cell culture medium. At 24 hours after transfection, cells were illuminated with red light using a custom-built 4×6 red light LED array (660 nm, Shenzhen Kiwi Lighting Co. Ltd; each LED centered above a single well). The light intensity was determined at the wavelength of 660 nm using an optical power meter (Q8230, Advantest), according to the manufacturer’s operating instructions. Reporter gene expression was quantified at 48 hours after illumination.

### Spatial control of REDMAP_Cre_ *in vitro*

HEK-293T cells (3.5×10^6^) were plated into a 10-cm dish, cultivated overnight to 60–70% confluency, and transfected with a total of 10 μg of plasmid DNA (pYZ208, pYZ247, and pDL78 at a ratio of 1:1:1 (w/w/w)). Twenty-three hours after the transfection, the cells were supplemented with 5 μM PCB in the cell culture medium. At 24 hours after transfection, the cells were placed on the display of a Vivo X9 smartphone with the pattern of a smiling face, and were illuminated with red light for 5 minutes (display brightness set to 100%, 40 μW cm^-2^). Fluorescence images of EGFP expression were acquired 24 hours after illumination using ChemiScope 4300 Pro imaging equipment (ChemiScope 4300Pro, Clinx).

### qPCR analysis

Total RNA was extracted from tissues employing an RNAiso Plus kit (Takara) according to the manufacturer’s instructions. Subsequently, the cDNA was synthesized using a HiScript^®^ II 1st Strand cDNA Synthesis Kit (+gDNA wiper) (R212-01, Vazyme) according to the manufacturer’s instructions. Real-time PCR was performed on a LightCycler 96 real-time PCR instrument (Roche) using the ChamQ Universal SYBR qPCR Master Mix (Q711, Vazyme) for target gene detection. The relative cycle threshold (CT) was determined and normalized to the mouse endogenous housekeeping gene *glyceraldehyde 3-phosphate dehydrogenase* (*Gapdh*). The results were expressed as a relative mRNA amount using the standard ^ΔΔ^Ct method.

### Immunoblotting analysis

Tissue samples were harvested and homogenized in RIPA lysis buffer (20101ES60, Yeasen) supplemented with protease inhibitor Cocktail (P1005, Beyotime). The resulting proteins were loaded onto SDS-PAGE gels and transferred to polyvinylidene difluoride membranes (PVDF; ISEQ00010, Millipore). The membranes were blocked with 5% non-fat milk for 2 hours at room temperature. The membranes were then incubated with anti-tdTomato primary antibody (A00682, GenScript) or anti-GAPDH primary antibody (10494-1-AP, Proteintech) overnight at 4 °C. After washing, the membranes were incubated with fluorochrome-conjugated secondary antibodies (33119ES60, Yeasen) at room temperature for one hour. Images were scanned using a fluorescent Western blot imaging system (LI-COR Odyssey Clx).

### AAV production

Adeno-associated viral vector (AAV8) carrying the *loxP*-STOP-*loxP*-Luciferase reporter was produced using previously described procedures^54^. Briefly, the packaging plasmid (pRC-2/8), helper plasmid (pHelper), and transfer plasmid (pAAV-Luci) were mixed at a 1:1:1 ratio and transfected into HEK-293FT cells. Seventy-two hours post-transfection, the cells were harvested and resuspended in lysis buffer, followed by three cycles of freezing and thawing. The AAV was then purified using iodixanol density gradient centrifugation and subsequently titrated using quantitative PCR. AAV8 and AAV9 that package the ΔPHYA-L5-CreC expression vector (pYZ590, ITR-P_hCMV_-ΔPhyA-CreC-pA-ITR), the FHY1-L3-CreN expression vector (pYZ591, ITR-P_hCMV_-FHY1-CreN-pA-ITR), and the all-in-one REDMAP_Cre_ expression vector (pYZ751, ITR-P_hCMV_-miniFHY1-CreN-P2A-ΔPhyA-CreC-pA-ITR) were produced by Packgene Biotech and applied iodixanol density gradient centrifugation for purification. Purified AAVs were titrated using quantitative PCR and concentrated in PBS to ∼1×10^13^ vg mL^-1^ (vg, vector genome).

### Animals

Six-week-old C57BL/6 male mice were obtained from the ECNU Laboratory Animal Centre. The *R26-tdTomato* transgenic mouse line [Ai14, *Gt(ROSA)26Sor^tm^*^14^*^(CAG-tdTomato)Hze^*]^34^ and the Rosa-LSL-UHRF1 mouse line^39^ have been described previously. The Rosa-LSL-DTA mouse line was obtained from Shanghai Model Organisms Center (SMOC). The REDMAP_Cre_ mouse line was generated by the Shanghai Model Organisms Center (SMOC). Briefly, the REDMAP_Cre_ transgenic mouse line was generated by inserting *P_CAG_-ΔPhyA-CreC-* WPRE-pA*-P_CAG_-FHY1-CreN-*WPRE-pA into the *Rosa26* locus using CRISPR-Cas9 (gRNA, 5’-GGGGACACACTAAGGGAGCTTGG-3’). REDMAP_Cre_ mice were crossed with Ai14, Rosa-LSL-UHRF1, or Rosa-LSL-DTA mouse lines to generate the corresponding double-positive offspring (REDMAP_Cre_:Ai14, REDMAP_Cre_:Rosa-LSL-UHRF1, and REDMAP_Cre_:Rosa-LSL-DTA). All progeny were genotyped by PCR to confirm the presence of the targeted alleles. All mice were kept on a standard alternating 12-h light/12-h dark cycle and given a normal chow diet [6% fat and 18% protein (w/w)] and water.

### Genomic PCR

Genomic DNA was prepared from the tails of mice. Tissues were lysed by incubation with lysis buffer (100 mM Tris HCl (pH 7.8), 5 mM EDTA, 0.2% SDS, 200 mM NaCl, and 100 μg mL^-1^ proteinase K) overnight at 55 °C, followed by centrifugation at 15,000 rpm for 5 minutes to obtain supernatants with genomic DNA. DNA was precipitated by isopropanol, washed in 70% ethanol, and dissolved in deionized water.

### Isolation and culture of primary cells

For fibroblast isolation, the skin from P1 to P3 mice was minced and digested with 0.125% trypsin/ethylenediaminetetraacetic acid (EDTA) solution (25200-072, Gibco). The cells were cultured in DMEM containing 10% (v/v) FBS and 1% (v/v) penicillin/streptomycin solution. For bone marrow-derived stem cells (BMSC) isolation, the bone marrow of the femurs of 6–8-week-old mice was flushed out with a culture medium. The cells were cultured in MEM Alpha Modification (SH30265.01, Cytiva) containing 10% (v/v) FBS (16000-044, Gibco) and 1% (v/v) penicillin/streptomycin solution. After 72 hours, nonadherent cells were removed, and fresh medium was added. For cortical cell isolation, the cells were isolated from Pl to P3 mice by using Papain (1495005, Sigma). The cells were harvested on Poly-L-lysine-coated dish with Neurobasal medium (10888022, Thermo-Fisher), 10% (v/v) FBS (16000-044, Gibco), 2% (v/v) B27 (A3582801, Thermo-Fisher), 1% (v/v) GlutaMax (35050061, Thermo-Fisher), and 1% (v/v) penicillin/streptomycin solution. The following day, the medium was changed to fresh medium without FBS.

### *In vivo* bioluminescence and imaging

Each mouse was intraperitoneally injected with D-Luciferin solution (150 mg kg^-1^; luc001, Shanghai Sciencelight Biology Science & Technology) and anesthetized with 2 % isoflurane (R510-22-10, RWD Life Science) dissolved in oxygen using an economical animal anesthesia machine (HSIV-S). Five minutes after luciferin injection, bioluminescence images of the mice were obtained using an IVIS Lumina II i*n vivo* imaging system (PerkinElmer), and bioluminescence images were analyzed with Living Image software (version 4.3.1).

### Histology

The mice were sacrificed, and fresh tissues were harvested and washed three times with cold PBS to remove impurities such as blood and then fixed in 4% (w/v) paraformaldehyde (PFA, 30525-89-4, Sangon Biotech) for 24 hours at 4 °C. Tissue blocks of ∼1 cm^3^ were cut and embedded in optimum cutting temperature compound (OCT, 03803389, Leica). Five µm thick tissue sections were prepared using Cryostat Microtome (CM1950, Leica) and rinsed with PBS. Finally, samples were counterstained with 4′,6-diamidino-2-phenylindole (DAPI, 28718-90-3, Sigma) for 10 minutes. The tdTomato signal was observed on an inverted fluorescence microscope (DMI8, Leica). Some slides were scanned using Pannoramic SCAN system (Pannoramic MIDI, 3DHISTECH).

### DNA recombination with REDMAP_Cre_ in C57BL/6 mice

Two milliliters (10% of the body weight in grams) of Ringer’s solution containing a total of 300 μg plasmids [100 µg pYZ208 (P_hCMV_-ΔPHYA-L5-CreC-pA), 100 µg pYZ247 (P_hCMV_-FHY1-L3-CreN-pA) and 100 µg pXY185 (P_hCMV_-*loxP*-STOP-*loxP*-Luciferase-pA)] was hydrodynamically injected into mice (male, six weeks old) via tail vein injection. At 8 hours after plasmid injection, the mice were intraperitoneally injected with 0 or 5 mg kg^-1^ PCB and then exposed to red light (660 nm, 20 mW cm^−2^) for different periods. Eight hours after initial illumination, bioluminescence images of the mice were obtained using an *IVIS* Lumina II *in vivo* imaging system (Perkin Elmer), and analyzed with the Living Image software (version 4.3.1). The mice were illuminated with an integrated LED (50 W, 45miL, with the light angle 60°, (660 ± 10) nm, Shenzhen Kiwi Lighting Co., Ltd) as described previously^33^.

### DNA recombination with REDMAP_Cre_ in Ai14 tdTomato transgenic reporter mice

Two milliliters (10% of the body weight in grams) of Ringer’s solution containing a total of 200 μg plasmids (100 µg pYZ208 and 100 µg pYZ247) were hydrodynamically injected into mice (male, six weeks old) via tail vein injection. At 12 hours after plasmid injection, the mice were intraperitoneally injected with 0 or 5 mg kg^-1^ PCB and then exposed to red light (660 nm, 20 mW cm^−2^) for one second or with no exposure. Mice were sacrificed, and livers were analyzed by qPCR, immunoblotting, and histology seven days after illumination.

For the AAV-REDMAP_Cre_-mediated DNA recombination in mouse livers, the Ai14 tdTomato reporter mouse (male, six weeks old) were tail vein injected with a mixture of 100 μL AAV8 virus containing two AAV vectors: the ΔPHYA-L5-CreC expression vector (pYZ590, ITR-P_hCMV_-ΔPhyA-CreC-pA-ITR, 2.5×10^11^ vg) and the FHY1-L3-CreN expression vector (pYZ591, ITR-P_hCMV_-FHY1-CreN-pA-ITR, 2.5×10^11^ vg). For the AAV-REDMAP_Cre_-mediated DNA recombination in mouse muscles, the transgenic mice (male, six weeks old) were intramuscularly injected into the left gastrocnemius muscle using an Omnican^®^ 40 (0.3 mm × 8 mm) syringe with a mixture of 50 μL AAV9 virus (1.5×10^11^ vg pYZ590 and 1.5×10^11^ vg pYZ591). For the single AAV-REDMAP_Cre_-mediated DNA recombination in mouse muscles, the transgenic mice (male, six weeks old) were intramuscularly injected into the left gastrocnemius muscle with 50 μL AAV9 virus (pYZ751, ITR-P_hCMV_-miniFHY1-CreN-P2A-ΔPhyA-CreC-pA-ITR, 5×10^11^ vg). For the single AAV-REDMAP_Cre_-mediated DNA recombination in mouse livers, the transgenic mice (male, six weeks old) were tail vein injected with 50 μL AAV8 virus (pYZ751, 5×10^11^ vg). Two weeks after injection, the transgenic mice were intraperitoneally injected with PCB (0 or 20 mg kg^-1^) and illuminated with red light (660 nm, 20 mW cm^-2^) for one hour or with no exposure. The mice were sacrificed, and livers and muscles were analyzed seven days after illumination.

### Characterizing genetic recombination activity in REDMAP_Cre_ mouse liver

The REDMAP_Cre_ transgenic mice (six weeks old) were tail vein injected with a mixture of 100 μL AAV8 virus carrying the *loxP*-STOP-*loxP*-Luciferase reporter (1.5×10^11^ vg). Two weeks after the AAV injection, the REDMAP_Cre_ transgenic mice were intraperitoneally injected with PCB (0 or 20 mg kg^-1^) and then exposed to red light (660 nm, 20 mW cm^-2^) or with no exposure. Bioluminescence was quantified one week after illumination using IVIS.

### Light-dependent DNA recombination in primary cells isolated from the REDMAP_Cre_:Ai14 mice

Mouse fibroblasts, bone marrow-derived stem cells (BMSCs), and cortical cells isolated from the REDMAP_Cre_:Ai14 mice were plated in a 24-well plate. The primer cells were supplemented with PCB (0 or 15 μM), and then illuminated with red light (660 nm, 1 mW cm^-2^) or with no exposure, tdTomato signals were profiled by fluorescence microscopy (Olympus IX71, TH4-200, Olympus) after illumination. Fluorescence intensity measurements were analyzed using Image J software.

### Light-dependent DNA recombination in REDMAP_Cre_:Ai14 mice

For DNA recombination in the livers, kidneys, muscles, hearts, lungs, intestines, spleens, and skin of the REDMAP_Cre_:Ai14 mice, six-week-old male mice were intraperitoneally injected with PCB (0 or 200 mg kg^-1^) and then illuminated with red light (660 nm, 20 mW cm^-2^) for one hour, or with no exposure. The mice were sacrificed, and the tissues were harvested and dissected for analysis seven days after illumination. For DNA recombination in the brains of the REDMAP_Cre_:Ai14 mice, six-week-old male mice were intracerebrally injected with 3 μL PCB (50 mg mL^-1^) using a 5.0-μL Neuros syringe (Hamilton) and then illuminated with red light (660 nm, 20 mW cm^-2^) for one hour. The mice were sacrificed, and the brains were isolated and sectioned at seven days after illumination. The tdTomato signal was observed on an inverted fluorescence microscope (DMI8, Leica).

### Spatial control of DNA recombination in REDMAP_Cre_:Ai14 mouse liver

The REDMAP_Cre_:Ai14 mice (male, six weeks old) were intraperitoneally injected with PCB (0 or 20 mg kg^-1^) and then locally irradiated using a laser pen (DL552004, Deli Group Co., Ltd.) for one minute (660 nm, 50 mW cm^-2^) from the abdomen. The mice were sacrificed, and the livers were harvested and dissected seven days after illumination. The tdTomato signal was scanned using a Pannoramic SCAN system (Pannoramic MIDI, 3DHISTECH).

### Light-dependent metabolic dysregulation in REDMAP_Cre_:Rosa-LSL-UHRF1 mice

REDMAP_Cre_:Rosa-LSL-UHRF1 transgenic mice were generated by crossing REDMAP_Cre_ mice with Rosa-LSL-UHRF1 mice, followed by genotypic screening to identify positive offspring. To investigate light-dependent metabolic dysregulation, five-week-old male REDMAP_Cre_:Rosa-LSL-UHRF1 mice were intraperitoneally injected with PCB (20 mg kg^-1^) and then illuminated with red light (660 nm, 20 mW cm^-2^) for one hour from the abdomen. One week post-treatment, glucose tolerance test (GTT) and insulin tolerance tests (ITT) were conducted to assess metabolic function. At the experimental endpoint, blood samples were collected for serum triglyceride (TG) analysis using a commercial assay kit (A110-1-1, Nanjing Jiancheng). Liver tissues were surgically excised, and Oil Red O staining was performed following standard protocols to evaluate lipid deposition.

### Glucose and insulin tolerance tests

For glucose tolerance tests (GTT), mice were fasted for 16 h, followed by intraperitoneal (i.p.) glucose injection (1.5 g/kg). For insulin tolerance tests (ITT), mice received an i.p. injection of insulin (1 U/kg). Blood glucose levels were monitored via tail vein blood samples at 0, 30, 60, 90 and 120 min after administration using a Contour Glucometer (Exactive Easy III, MicroTech Medical). The trapezoidal rule was used to determine the area under the curve (AUC) for GTT and ITT.

### Light-dependent cell ablation in REDMAP_Cre_:Rosa-LSL-DTA mice

REDMAP_Cre_:Rosa-LSL-DTA transgenic mice were generated by crossing REDMAP_Cre_ mice with Rosa-LSL-DTA mice, followed by genotypic screening to identify positive offspring. To investigate light-dependent cell ablation, six-week-old REDMAP_Cre_:Rosa-LSL-DTA mice were intraperitoneally injected with PCB (20 mg kg^-1^) and then illuminated with red light (660 nm, 20 mW cm^-2^) for one hour from the abdomen. Twenty-four hours post-treatment, the mice were humanely euthanized, and serum samples were collected for assess hepatic injury. Levels of the liver injury biomarkers, alanine aminotransferase (ALT) and aspartate aminotransferase (AST), were quantified using commercial assay kits (ALT: C009-2-1; AST: C010-2-1; Nanjing Jiancheng), following the manufacturer’s protocols.

### Ethics

The experiments involving animals were approved by the East China Normal University (ECNU) Animal Care and Use Committee and were in direct accordance with the Ministry of Science and Technology of the People’s Republic of China on Animal Care guidelines. The protocol (protocol ID: m20230205) was approved by the ECNU Animal Care and Use Committee. All mice were euthanized after the experiments.

### Quantification and statistical analysis

Unless otherwise mentioned, all *in vitro* data represent the mean ± SD of three independent biological replicates. For the animal experiments, each group consisted of randomly selected mice (*n* = 3-5), and the results are expressed as mean ± SEM. Comparisons between groups were performed using Student’s *t*-tests. One-way ANOVA was used to evaluate differences between multiple groups with a single intervention, followed by the Dunnett post hoc test. Statistical significance was determined using GraphPad (Prism v.8), and the significance was assigned at \**P* < 0.05, \*\**P* < 0.01 and \*\*\**P* < 0.001. *n* and *P* values are described in the figures or figure legends.

## Data availability

All data associated with this study are present in the paper or the Supplementary Information. All genetic components related to this paper are available with a material transfer agreement and can be requested from H.Y. Source data are provided in this paper.

## Acknowledgments

This work was financially supported by grants from the National Natural Science Foundation of China (no. 31861143016 and 31971346), the National Key R&D Program of China, Synthetic Biology Research (no. 2019YFA0904500), the Science and Technology Commission of Shanghai Municipality (no. 22N31900300 and 18JC1411000), and the Fundamental Research Funds for the Central Universities to H.Y. This work was also partially supported by the Young Scientists Fund of the National Natural Science Foundation of China (no. 32300458), the Science and Technology Commission of Shanghai Municipality (no. 23YF1410700), China Postdoctoral Science Foundation (no. 2022M721163 and no. BX20230128), and the Natural Science Foundation of Chongqing (CSTB2023NSCQ-MSX0126) to Y.Z, as well as the National Natural Science Foundation of China (no. 32301217) to J.Y and Natural Science Foundation of Shandong Province (no. ZR2021LLZ003) to W.L. We also thank the support from the CAS Youth Interdisciplinary Team, and the ECNU Multifunctional Platform for Innovation (011), and the Instruments Sharing Platform of the School of Life Sciences, ECNU regarding the mice experiments.

## Author contributions

H.Y. conceived the project. H.Y., Y.Z., and Y.W. designed the experiments, analyzed the results, and wrote the manuscript. Y.Z., Y.W., J.Y., D.K., W.L., X.W., Y.Y., G.Y., L.L., L.Q., and H.L. performed the detailed experiments. H.Y., Y.Z., Y.W., J.Y., D.L., and N.G. analyzed the results and wrote the manuscript. All authors revised and approved the manuscript.

