## Supplementary information for "A rapid and efficient red-light-activated Cre recombinase system for genome engineering in mammalian cells and transgenic mice"

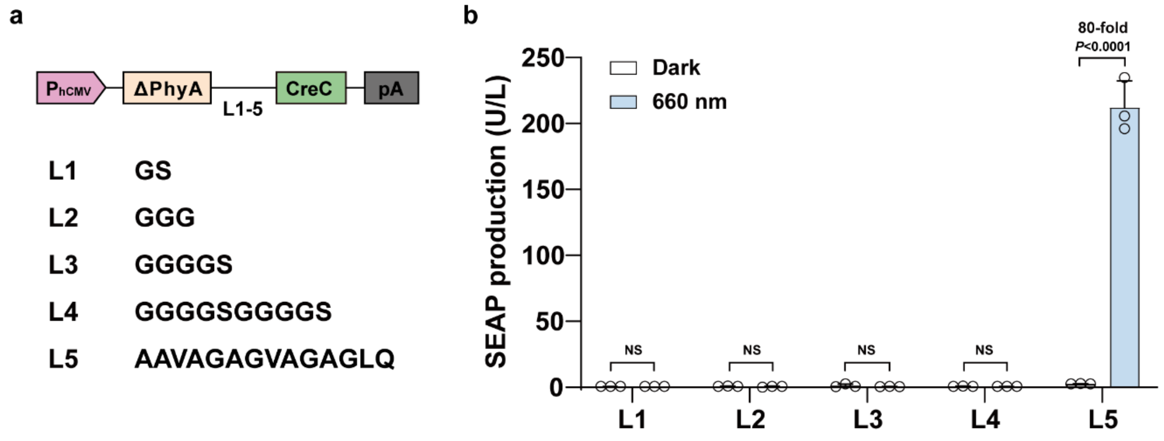


**Supplementary Fig. 1 | Optimization for recombination efficiency using the different linkers (L1 to L5) between the ΔPhyA and CreC domains.** HEK-293T cells ($6\times10^4$) co-transfected with FHY1-L3-CreN expression vector (pYZ247), the Cre-dependent SEAP reporter (pGY125), and ΔPhyA-CreC fusion protein expression vector (pYZ245/pYZ246/pYZ247/pYZ248/pYZ209) with different linkers (L1 to L5) were supplied with PCB (5 μM) and then illuminated with red light (660 nm, 1 mW $cm^{-2}$) for 48 hours; SEAP production in the culture supernatant was quantified after illumination. All data are presented as means ± SD. Student's *t*-tests were used for comparison. *n* = 3 independent experiments. NS, not significant.

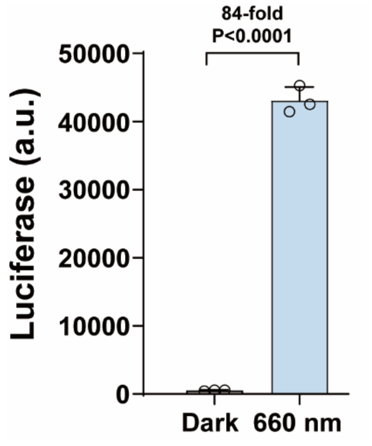


**Supplementary Fig. 2 | Cre-catalyzed DNA recombination with $REDMAP_{Cre}$ using a luciferase**

**reporter.** HEK-293T cells ($6\times10^4$) cells co-transfected with ΔPhyA-L5-CreC (pYZ208), FHY1-L3-CreN (pYZ247), and the Cre-dependent Luciferase reporter (pXY185, $P_{hCMV}$-*loxP*-STOP-*loxP*-Luciferase-pA) were supplied with PCB (5 μM) and then illuminated with red light (660 nm, 1 mW cm$^{-2}$) for 48 hours; Bioluminescence measurements were taken after illumination. All data are presented as means ± SD. Student's *t*-tests were used for comparison. *n* = 3 independent experiments.

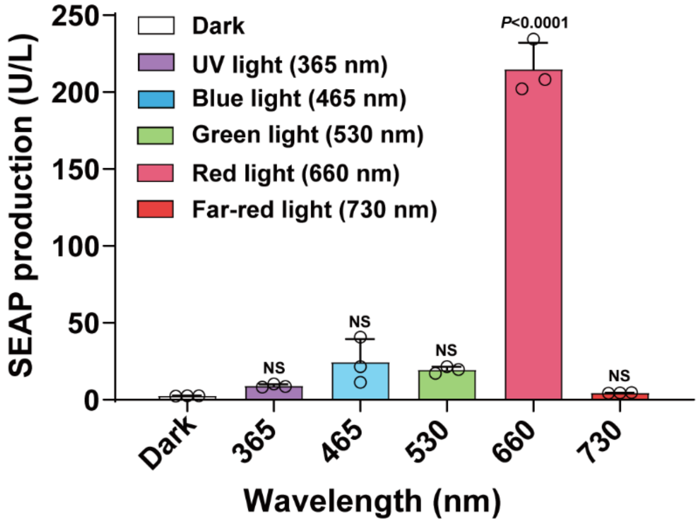


**Supplementary Fig. 3 | Chromatic specificity of REDMAP$_{\mathbf{Cre}}$.** HEK-293T cells ($6\times10^4$) co-transfected with ΔPhyA-L5-CreC (pYZ208), FHY1-L3-CreN (pYZ247), and the Cre-dependent SEAP reporter (pGY125) were supplied with PCB (5 μM) and then illuminated with different wavelengths of light (from 365 nm to 730 nm as indicated, 1 mW cm$^{-2}$) for one minute; SEAP production in the culture supernatant was quantified 48 hours after illumination. All data are presented as means ± SD. One-way ANOVA was used for comparison. *n* = 3 independent experiments. NS, not significant.

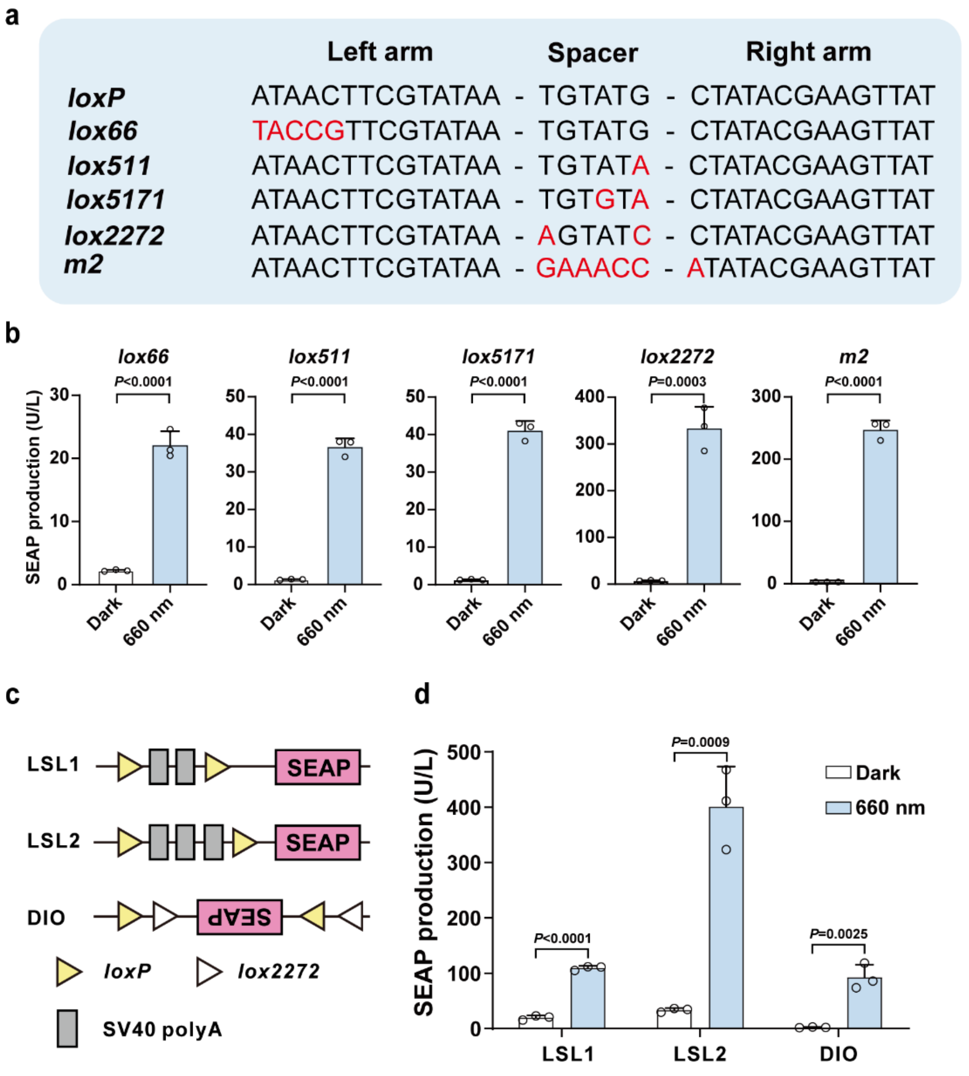


**Supplementary Fig. 4 | REDMAP$_{Cre}$-catalyzed DNA recombination capacity with different Cre recombinase-dependent reporters.** **a**, Different mutated *loxP* sequences and mutations are highlighted in red. **b**, The catalytic recombination activity of REDMAP$_{Cre}$ for different *loxP* mutants. HEK-293T cells (6×10$^4$) co-transfected with ΔPhyA-L5-CreC (pYZ208), FHY1-L3-CreN (pYZ247), and different plasmids encoding *loxP* mutants (pXY230/pWY152/pWY174/pXY229/pWY175) were supplied with PCB (5 μM) and then illuminated with red light (660 nm, 1 mW cm$^{-2}$) for 48 hours; SEAP production in the culture supernatant was quantified after illumination. All data represent the mean ± SD; *n* = 3 independent experiments. **c**, Different constructions of Cre-dependent reporters. LSL1, two copies of the SV40 poly (A) structure between *loxP* sites; LSL2, three copies of the SV40 polyA structure between *loxP* sites; DIO, Cre-dependent double-floxed inverted open reading frame (DIO) expressing SEAP reporter. The SEAP reporter is expressed after two rounds of recombination at the *loxP* sites. **d**, The recombination activity of $REDMAP_{Cre}$ in different reporters. HEK-293T cells ($6\times10^4$) were co-transfected with ΔPhyA-L5-CreC (pYZ208), FHY1-L3-CreN (pYZ247), and different reporter plasmids (pDQ584/pWY78/pWY79). Then, the transfected cells were supplied with PCB (5 μM) and illuminated with red light (660 nm, 1 mW $cm^{-2}$) for 48 hours. SEAP production was quantified after illumination. All data are presented as means ± SD. Student's *t*-tests were used for comparison. *n* = 3 independent experiments.

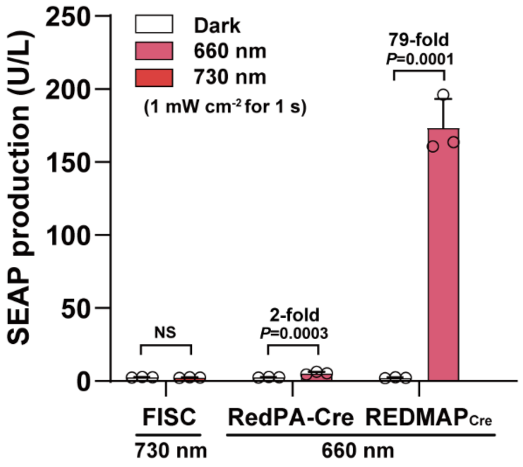


**Supplementary Fig. 5 | Comparison of $REDMAP_{Cre}$ with other red/far-red light-inducible recombinase systems (FISC and RedPA-Cre).** HEK-293T cells ($6\times10^4$) co-transfected with the Cre-dependent SEAP reporter (pGY125) and $REDMAP_{Cre}$ (pYZ208/pYZ247) or the FISC (pXY137/pXY237) or RedPA-Cre (pWY105/pWY107) were supplied with PCB (5 μM) and then illuminated with red (660 nm, 1 mW $cm^{-2}$) or far-red light (730 nm, 1 mW $cm^{-2}$) for one second; SEAP production in the culture supernatant was quantified 48 hours after illumination. All data are presented as means ± SD. Student's *t*-tests were used for comparison. *n* = 3 independent experiments. NS, not significant.

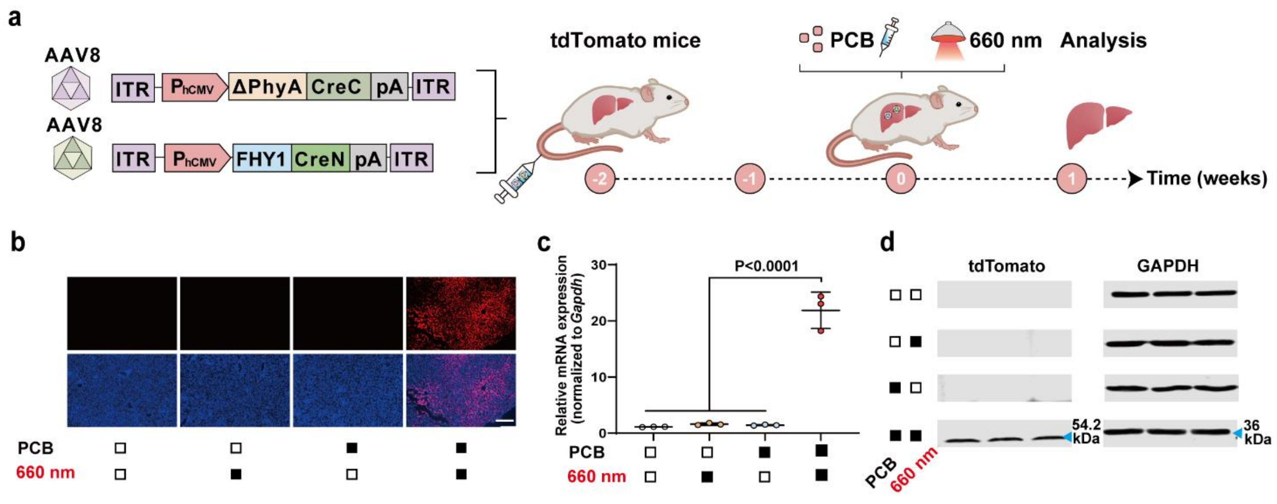


**Supplementary Fig. 6 | AAV-REDMAP$_{Cre}$-mediated DNA recombination in mouse livers. a**, Schematic representation of the experimental procedure for AAV8 delivery of REDMAP$_{Cre}$ to Ai14 tdTomato reporter mouse livers. Mice were tail vein injected with a mixture of AAV encoding the REDMAP$_{Cre}$. After two weeks, mice were intraperitoneally injected with PCB (20 mg kg$^{-1}$) and illuminated with red light (660 nm, 20 mW cm$^{-2}$) for an hour. The control mice were exposed to either red light, PCB alone, or neither. The mice were sacrificed, and their livers were analyzed seven days after illumination. **b** Representative fluorescence images of liver sections from the indicated groups. Blue, DAPI. Red, tdTomato. Scale bar, 200 µm. **c-d**, qPCR (**c**), and immunoblotting (**d**) analysis of tdTomato in isolated liver tissues. Black block, with treatment; white block, without treatment. Data in **c** are expressed as means ± SEM. One-way ANOVA was used for multiple comparisons. *n* = 3 mice.

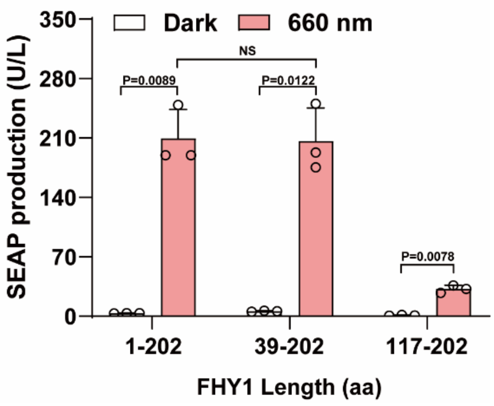

**Supplementary Fig. 7 | Minimizing and characterizing the different truncated FHY1 constructs fused to the CreN domain.** HEK-293T cells (6×10$^4$) co-transfected with the ΔPhyA-L5-CreC expression vector (pYZ208), the Cre-dependent SEAP reporter (pGY125), and different truncated versions of FHY1: pYZ247 [$P_{hCMV}$-FHY1(1-202aa)-L3-CreN-pA], pYZ744 [$P_{hCMV}$-FHY1 (117-202 aa)-L3-CreN-pA] or pYZ746 [$P_{hCMV}$-miniFHY1-L3-CreN-pA; miniFHY1, FHY1 (39-202 aa)]. Twenty-four hours after transfection, cells were supplied with PCB (5 μM) and then illuminated with red light (660 nm, 1 mW cm$^{-2}$) for one minute; SEAP production in the culture supernatant was quantified 48 hours after illumination. aa, amino acids. All data are presented as means ± SD. Student's *t*-tests were used for comparison. *n* = 3 independent experiments. NS, not significant.

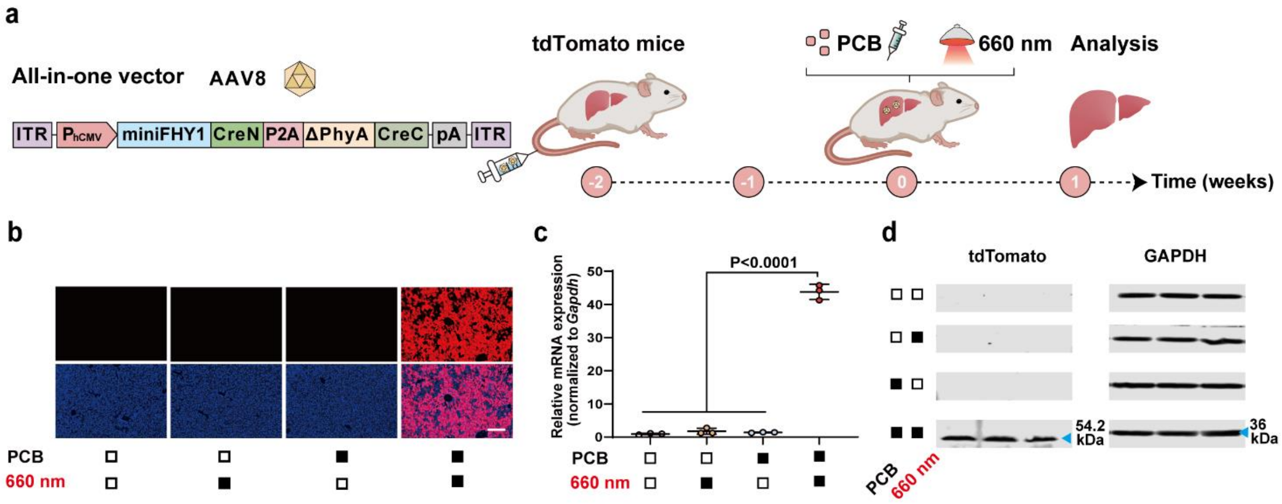


**Supplementary Fig. 8 | All-in-one AAV-REDMAP$_{Cre}$-mediated DNA recombination in mouse liver. a**, Schematic representation of the experimental procedure for AAV8 delivery of REDMAP$_{Cre}$ to Ai14 tdTomato reporter mouse livers. Mice were tail vein injected with an all-in-one AAV encoding the REDMAP$_{Cre}$ (pYZ751). After two weeks, mice were intraperitoneally injected with PCB (20 mg kg$^{-1}$) and illuminated with red light (660 nm, 20 mW cm$^{-2}$) for an hour. The control mice were exposed to either red light, PCB alone, or neither. The mice were sacrificed, and their livers were analyzed seven days after illumination. **b,** Representative fluorescence images of liver sections from the indicated groups. Blue, DAPI. Red, tdTomato. Scale bar, 200 µm. **c-d**, qPCR (**c**), and immunoblotting (**d**) analysis of tdTomato in isolated liver tissues. Black block, with treatment; white block, without treatment. Data in **c** are expressed as means ± SEM. One-way ANOVA was used for multiple comparisons. *n* = 3 mice.

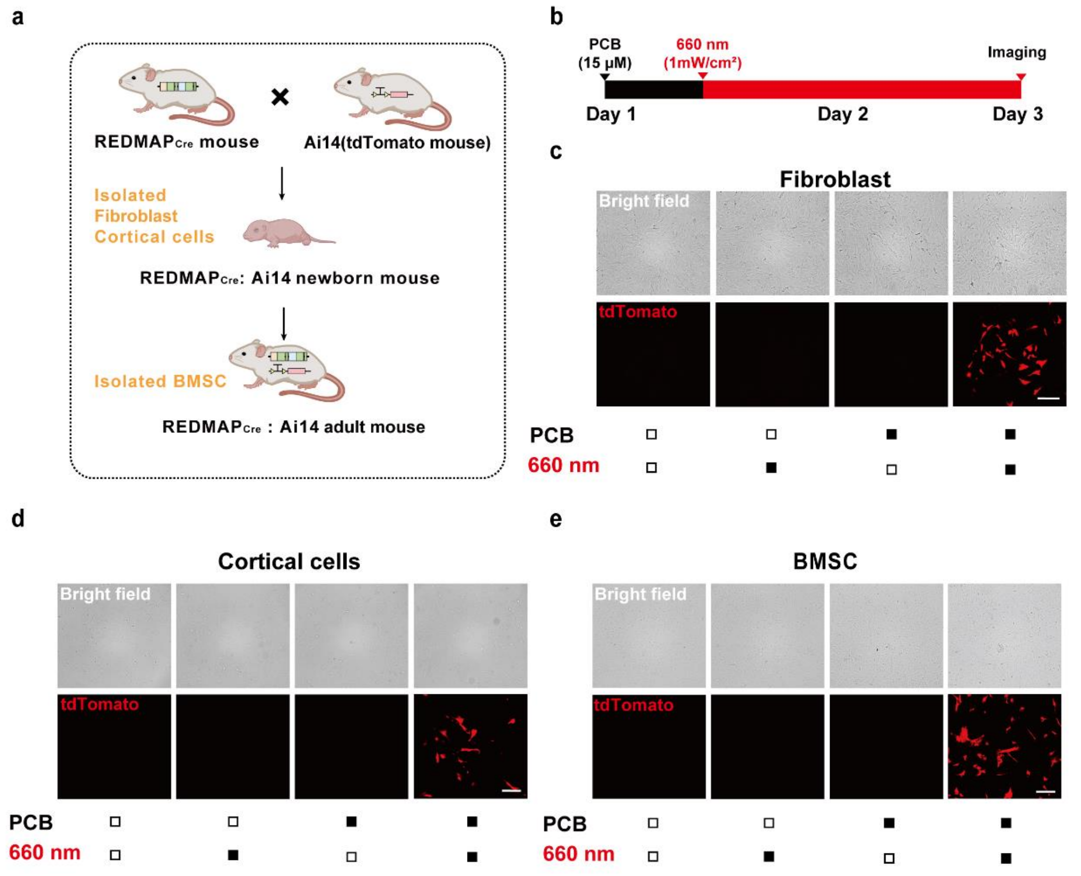


**Supplementary Fig. 9 | Light-dependent recombinase activity in primary cells isolated from the REDMAP$_{Cre}$:Ai14 mice. a,** A schematic diagram illustrates the hybridization of REDMAP$_{Cre}$ mice and Ai14 mice. The offspring of these mice, REDMAP$_{Cre}$ mice, were used for subsequent experiments as double-heterozygous subjects. Fibroblasts and cortical cells were obtained from newborn REDMAP$_{Cre}$ mice, while bone marrow-derived stem cells (BMSCs) were obtained from adult mice. **b,** Schematic diagram for evaluating light-dependent recombinase activity in primary cells from the

REDMAP$_{Cre}$:Ai14 mice. Fibroblasts, cortical cells and BMSC were supplemented with 0 or 15 μM PCB, then illuminated with red light (660 nm, 1 mW cm$^{-2}$) for 48 hours, tdTomato levels were profiled by fluorescence microscopy after illumination. **c-e**, Representative tdTomato fluorescence images of fibroblast (**c**), cortical cells (**d**), BMSC (**e**) from REDMAP$_{Cre}$:Ai14 mice. Black block, with treatment; white block, without treatment. Top images, the bright field, bottom images, the tdTomato field. Scale bar, 200 μm.

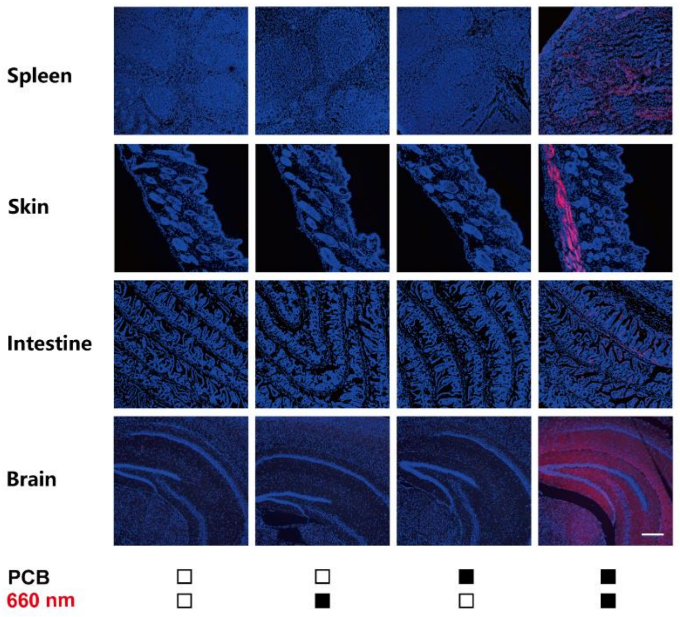


**Supplementary Fig. 10 | Fluorescence images of various organ sections from REDMAP$_{Cre}$:Ai14 mice.** Adult REDMAP$_{Cre}$:Ai14 mice were intraperitoneally injected with/without PCB (200 mg kg$^{-1}$) and then exposed to red light (660 nm, 20 mW cm$^{-2}$) for one hour. The mice were sacrificed, and their spleen, skin, and intestines were harvested and analyzed seven days after the illumination. Black block, with treatment; white block, without treatment. Blue, DAPI. Red, tdTomato. Scale bar, 200 µm.
